# The multi PAM2 protein Upa2 functions as novel core component of endosomal mRNA transport

**DOI:** 10.1101/455378

**Authors:** Silke Jankowski, Thomas Pohlmann, Sebastian Baumann, Kira Müntjes, Senthil Kumar Devan, Sabrina Zander, Michael Feldbrügge

**Author notes:** equal contribution (Silke Jankowski, Thomas Pohlmann). Cell and Developmental Biology, Centre for Genomic Regulation (CRG), Barcelona, Spain. Corresponding author: Dr. Michael Feldbrügge, Institute for Microbiology, Cluster of Excellence on Plant Sciences, Heinrich Heine University Düsseldorf; 40204 Düsseldorf, Germany.

## Abstract

mRNA transport determines spatiotemporal protein expression. Transport units are higher-order ribonucleoprotein complexes containing cargo mRNAs, RNA-binding proteins and accessory proteins. Endosomal mRNA transport in fungal hyphae belongs to the best-studied translocation mechanisms. Although several factors are known, additional core components are missing. Here, we describe the 232 kDa protein Upa2 containing multiple PAM2 motifs (poly[A]-binding protein [Pab1] associated motif 2) as a novel core component. Loss of Upa2 disturbs transport of cargo mRNAs and associated Pab1. Upa2 is present on almost all transport endosomes in an mRNA dependent-manner. Surprisingly, all four PAM2 motifs are dispensable for function during unipolar hyphal growth. Instead, Upa2 harbours a novel N-terminal effector domain as important functional determinant as well as a C-terminal GWW motif for specific endosomal localisation. In essence, Upa2 meets all the criteria of a novel core component of endosomal mRNA transport and appears to carry out crucial scaffolding functions.

## Introduction

Active transport of mRNAs determines when and where proteins are synthesised. Such trafficking events are important for a wide variety of different cellular processes like asymmetric cell division, polar growth, embryonic development and neuronal activity [1, 2]. Several mRNA translocation mechanisms have been described. During cytokinesis of *Saccharomyces cerevisiae*, for example, the actin-dependent transport of *ASH1* mRNA is mediated by the concerted binding of the RNA-binding proteins (RBP) She2p and She3p. These RBPs connect the cargo mRNA to the myosin motor Myo4 for transport towards the daughter cell [3, 4]. In highly polarised cells such as fungal hyphae and neurons, mRNAs are transported along microtubules over long distances. Here, molecular motors such as kinesins and dynein are involved. Among the best studied examples of microtubule-dependent translocation is the mRNA transport on shuttling endosomes during polar hyphal growth in the fungus *Ustilago maydis* [5].

Upon infection of corn, *U. maydis* switches from budding to hyphal growth [6, 7]. The resulting infectious hyphae grow with a defined axis of polarity: they expand at the apical pole and insert septa at the basal pole resulting in the formation of characteristic sections devoid of cytoplasm. In this growth mode, hyphae depend on active transport along microtubules. Loss of long-distance transport results in aberrant hyphal growth. Characteristic of this defect is the formation of bipolar growing cells. Important carriers are Rab5a-positive endosomes that shuttle along microtubules by the concerted action of the plus-end directed Kinesin-3 type motor Kin3 and the minus-end directed cytoplasmic dynein Dyn1/2 [8, 9]. These endosomes carry characteristic markers of early endosomes involved in endocytosis [10]. However, during polar growth they also function as transport endosomes moving organelles, like peroxisomes, and mRNAs with associated ribosomes attached to their cytoplasmic surface [11, 12].

The key RNA-binding protein for mRNA transport is Rrm4, an RRM (RNA recognition motif) protein containing tandem N-terminal RRMs separated by a spacer from a third RRM domain (Fig 1A). The mRNAs of all four septins were identified to be important cargo mRNAs, which are most likely translationally active while being transported on endosomes [12–14]. Consequently, the translation products Cdc3, Cdc10, Cdc11 and Cdc12 form heteromeric complexes on the cytoplasmic surface of endosomes in an Rrm4-dependent manner. These septin complexes are delivered to the hyphal growth pole to form a longitudinal gradient of filaments [12, 14]. A recent transcriptome-wide view of endosomal mRNA transport revealed that Rrm4 binds thousands of mRNAs preferentially in their 3′ UTR in close proximity to the small glycine rich protein Grp1. This extensive mRNA transport is most likely needed for the distribution of mRNAs within the fast-growing hypha as well as for the transport of specific mRNAs encoding e.g. septins for heterooligomerisation [15].

**Figure 1.**
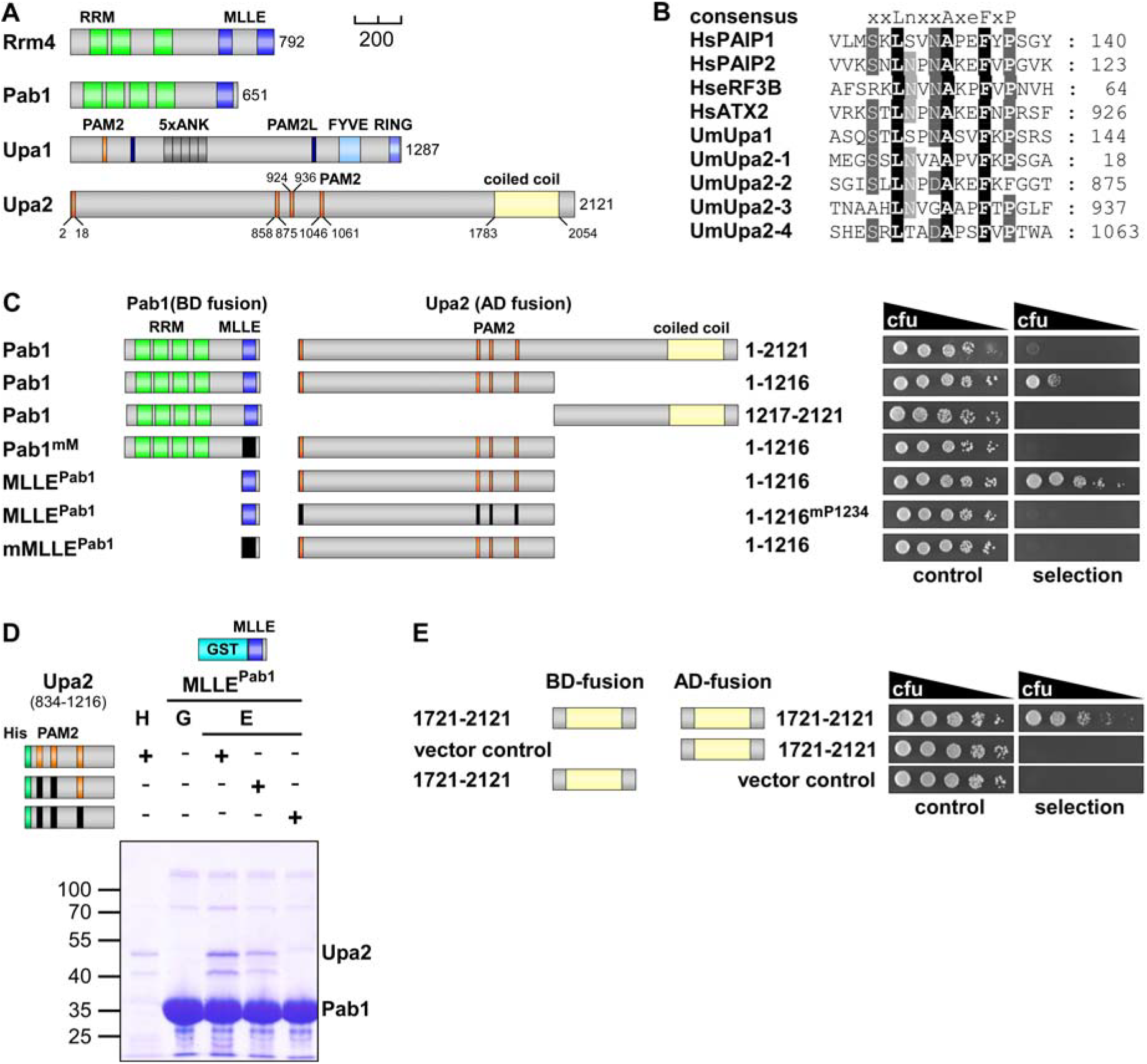
Upa2 contains multiple PAM2 motifs for interaction with Pab1. (**A**) Schematic representation of the MLLE domain-containing proteins Rrm4 and Pab1 as well as the PAM2 motif-containing proteins Upa1 and Upa2 drawn to scale (bar, 200 amino acids; green, RRM domain; blue, MLLE domain; orange, PAM2 motif; dark blue, PAM2-like motif; dark grey, Ankyrin repeats; light blue, FYVE domain; lilac, RING domain; yellow, coiled coil region) (**B**) Comparison of PAM2 sequences found in Upa2 (accession number UMAG_10350) with those of human proteins such as PAIP1 (accession number NP_006442.2), PAIP2 (accession number CAG38520.1), eRF3B (accession number CAB91089.1) and ATX2 (accession number NP_002964.3). (**C**) Two-hybrid analysis with schematic representation of protein variants tested on the left. Yeast cultures were serially diluted 1:5 (decreasing colony forming units, cfu) and spotted on respective growth plates controlling transformation and assaying reporter gene expression (see Materials and methods). (**D**) GST co-purification experiments with components expressed in *E. coli*: MLLE domain of Pab1 fused to GST and N-terminal His_6_-tagged versions of Upa2 (amino acids 834–1216; orange rectangles indicate PAM2 motif 1, 2 and 3; black rectangles mark mutations; 43 kDa). Interaction studies were performed with the soluble fraction of protein extracts from *E. coli* to demonstrate specific binding. After GST affinity chromatography proteins were eluted (lanes marked with “E”; G, input of GST-MLLE^Pab1^ and H, Ni-NTA precipitated His_6_-Upa2). Note that the Upa2 variants were enriched in the eluate due to the pull down by MLLE^Pab1^. (**E**) Two-hybrid analysis of the coiled coil region of Upa2 (amino acids 1712-2121) fused to binding domain (BD) and activation domain (AD) of the Gal4 transcription factor (design as in C).

At its C-terminus, Rrm4 carries two MademoiseLLE (MLLE) domains that function as peptide binding pockets for specific interaction with the PAM2-like motifs of Upa1. Upa1 links Rrm4-containing mRNPs to transport endosomes using a FYVE zinc finger domain to recognise phosphatidylinositol-3-phosphate lipids of early endosomes. [16]. In addition, Upa1 also carries a classical PAM2 motif for interaction with the MLLE domain of the poly(A)-binding protein Pab1, an additional component of endosomal mRNPs [7, 17]. Although we already identified a number of components of endosomal mRNA transport, key factors might still be missing. Here, we unravelled and characterized one such factor, the protein Upa2, a second *Ustilago* PAM2 motif protein that had previously been identified by bioinformatic prediction of PAM2-containing proteins [16].

## Results

### The PAM2-containing protein Upa2 interacts with Pab1

Upa2 (*Ustilago* PAM2 protein 2; UMAG_10350) is a 2121 amino acid (aa) protein with a conserved coiled coil domain of unknown function at its C-terminus, as well as four PAM2 motifs for potential interaction with the MLLE domain of Pab1. One PAM2 motif is situated at the immediate N-terminus and three in the central region of the protein (Fig 1A-B).

In order to validate the predicted PAM2 motifs we performed yeast two-hybrid studies that have already been successfully applied to demonstrate an interaction between Pab1 and the PAM2 motif of Upa1 [16]. Full-length Upa2 shows weak interaction with Pab1 but did not interact with Upa1 or Rrm4 (Fig 1C, Fig EV1A-B). The latter is consistent with the observation that PAM2-like motifs for Rrm4 interaction are missing in Upa2 [16]. Upon mapping the interaction domain of Upa2 with Pab1 we observed that the PAM2-containing N-terminal part of Upa2 (aa 1-1216) but not the C-terminal part (aa 1217-2121) interacted with Pab1 (Fig 1C). The interaction of Upa2 with Pab1 was mediated by the MLLE domain of Pab1, since this domain was necessary and sufficient for reporter gene expression in the yeast two-hybrid system (Fig 1C, Fig EV1A). Mutational analysis of the PAM2 motifs in Upa2 revealed that a single PAM2 motif was sufficient for interaction with the MLLE domain of Pab1 or full length Pab1 (Fig EV1A). Mutating all PAM2 motifs resulted in loss of interaction of Upa2 with the MLLE domain of Pab1 (Fig 1C) and strongly reduced interaction with Upa2 and full length Pab1 (Fig EV1A).

To verify this binding behaviour, we tested the interaction of the central PAM2 triplet of Upa2 (aa 834-1216) with the MLLE domains of Pab1 by GST-pulldown experiments using variants expressed in *Escherichia coli*. These experiments confirmed the specific interaction of the PAM2 motifs of Upa2 with the MLLE domain of Pab1. The interaction strength was dependent on the number of functional PAM2 motifs (Fig 1D). As expected, PAM2 motifs of Upa2 did not interact with Rrm4 (Fig EV1C-D).

Since coiled coil domains have been described as dimerization regions, we tested this in a yeast two-hybrid assay. When appended to both BD and AD reporter proteins the predicted C-terminal coiled coil domain of Upa2 supported yeast growth on selective medium suggesting that this region indeed functions as a dimerization domain (Fig 1E). Taken together, Upa2 contains multiple functional PAM2 motifs for interaction with Pab1 and a C-terminal dimerization domain. The interaction with Pab1 is the first hint that Upa2 might function in endosomal mRNA transport.

### Upa2 is essential for efficient unipolar hyphal growth

If Upa2 is indeed important for endosomal mRNA transport, loss of Upa2 should exhibit phenotypes similar to mutations in the previously identified transport components Rrm4 and Upa1 [16]. To test this assumption, we generated *upa2* deletion mutants in the genetic background of laboratory strain AB33. In this strain, hyphal growth can be elicited synchronously and reproducibly by switching the nitrogen source in the medium [18]. Hyphae grow with a defined axis of polarity: they expand at the apical tip and insert septa at their base leading to the formation of characteristic empty sections (Fig 2A). While loss of Upa2 did not affect growth or alter shape of yeast cells (Fig EV2A-B) *upa2*Δ strains exhibited a bipolar growth phenotype and the formation of empty sections was delayed. This aberrant growth mode is typical for defects in microtubule-dependent transport and has been described for loss of the key proteins of endosomal mRNA transport like Rrm4 and Upa1 (Fig 2A-B). Analysing the *upa1*Δ/*upa2*Δ double mutant revealed that the number of hyphae exhibiting aberrant bipolar hyphal growth was not additive (Fig EV2C), providing genetic evidence that both PAM2-containing proteins function in the same cellular process.

**Figure 2.**
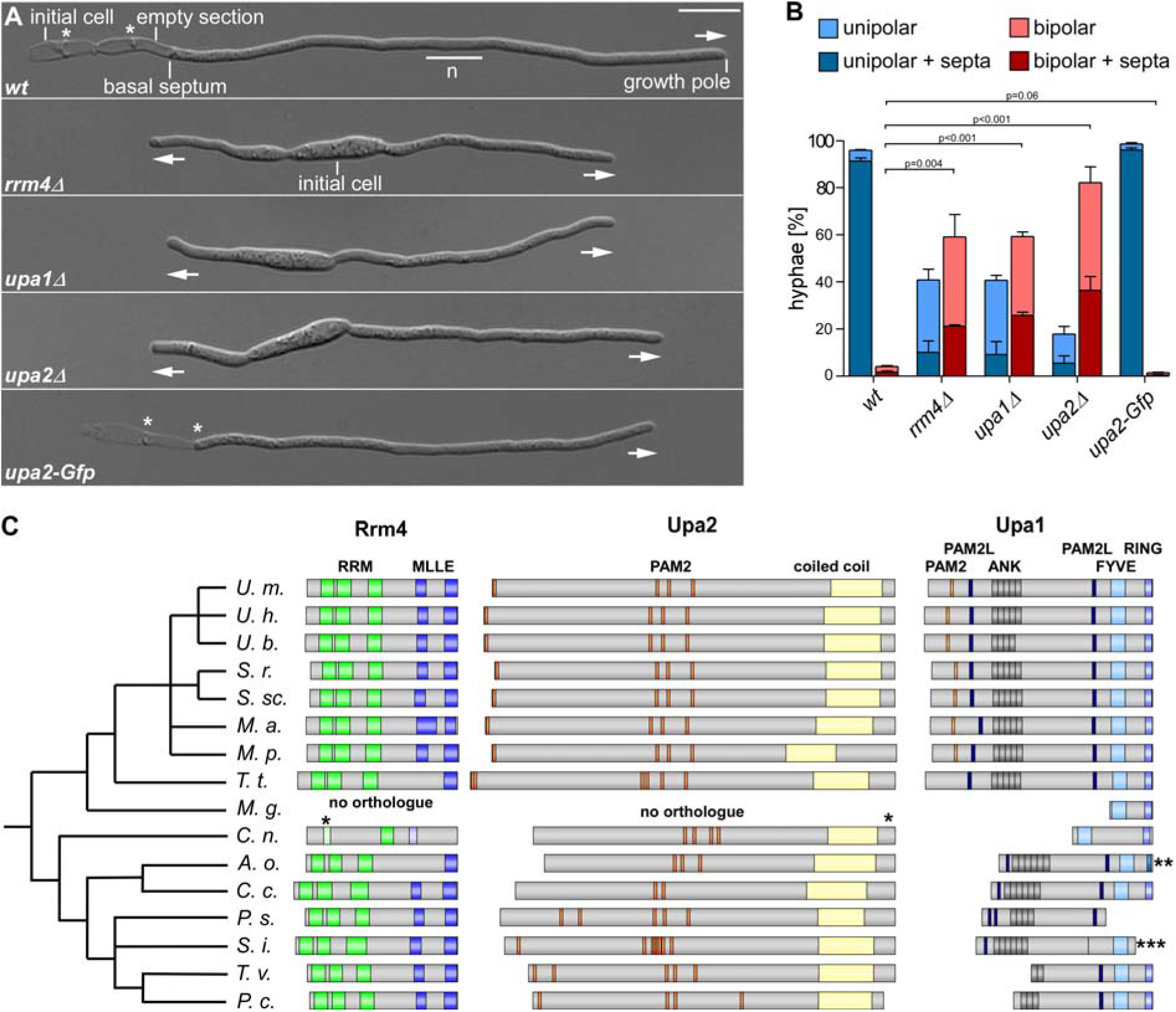
Loss of Upa2 causes defects in hyphal growth. (**A**) Growth of AB33 derivatives in the hyphal form (6 h.p.i.; growth direction is marked by arrows; n, nucleus; asterisks, basal septa). (**B**) Quantification of hyphal growth (6 h.p.i): unipolarity, bipolarity and septum formation were quantified (error bars, s.e.m.; n = 3 independent experiments, for each experiment >100 hyphae were counted per strain; note that septum formation is given relative to the values of unipolar or bipolar hyphae set to 100%). For statistical analysis, the percentage of bipolarity was investigated by using unpaired two-tailed Student′s t-test. Three independent experiments (n = 3) were conducted with at least 100 hyphae per strain. (**C**) Schematic representation of orthologues of Rrm4, Upa2 and Upa1 in different basidiomycetes. The phylogenetic tree was generated using the phyloT tool using the NCBI taxonomy (no evolutionary distances): U.m.: *Ustilago maydis*, U.h.: *Ustilago hordei*; U.b.: *Ustilago bromivora*; S.r.: *Sporisorium reilianum*; S.sc.: *Sporisorium scitamineum*, M.a.: *Moesziomyces antarcticus*, M.p.: *Melanopsichium pennsylvanicum*, T.t.: *Thecaphora thlaspeos*; M.g.: *Malassezia globosa*; C.n.: *Cryptococcus neoforman var neoformanss*; A.o.: *Armillaria ostoyae*; C.c.: *Coprinopsis cinerea*; P.s.: *Punctularia strigosozonata*; S.i.: *Serendipita indica*; T.v.: *Trametes versicolor*; P.c.: *Phanerochaete carnosa*. * Bioinformatic tools predict weak RRM domains and a non-functional MLLE domain in C.n. Rrm4. ** A.o. Upa1 contains a predicted transmembrane domain instead of a RING domain; *** S.i. Upa1 was manually assembled from two consecutive ORFs. Accession numbers are listed in Appendix Table S7.

A second read-out for defects in the endosomal mRNA transport machinery is the unconventional secretion of chitinase Cts1 [16, 19]. Loss of Upa2 resulted in reduced extracellular Cts1 activity specifically during hyphal growth. This is reminiscent of defective Cts1 secretion upon deletion of *rrm4* and *upa1* (Fig EV2D). Hence, Upa2 might function in concert with Rrm4 and Upa1 during endosomal mRNA transport.

This hypothesis was supported by a phylogenetic analysis showing that orthologs of Upa2 were found in several basidiomycete fungi with comparable spacing of the predicted PAM2 motifs (Fig 2C). Consistent with the notion that Rrm4, Upa1 and Upa2 function together, these fungi also had orthologs for Rrm4 and Upa1. Notably, Upa2 was absent in *Malassezia globosa*, which presumably has lost the endosomal mRNA transport machinery, because it lacks a clear ortholog of Rrm4 (Fig 2C) [20]. In essence, the evolutionarily conserved Upa2 is a novel factor essential for efficient unipolar growth that might function in endosomal mRNA transport.

### Upa2 shuttles on Rrm4-positive transport endosomes

*upa2* and *rrm4* appear to be genetically and phylogenetically linked. We therefore examined whether the protein can also be found on transport endosomes by expressing a functional version of Upa2 with C-terminally fused Gfp (eGfp, Clontech; Fig 2A-B; Fig EV2D). Fluorescence microscopy revealed that in hyphae Upa2-Gfp localised exclusively in bidirectionally moving units (Fig 3A, Video EV1). Movement was inhibited by benomyl indicating that it was microtubule-dependent (Fig 3A, Video EV1). Importantly, Upa2-Gfp shuttling resembled the movement of Rrm4-Gfp and Pab1-Gfp (Fig 3A, Video EV1). This holds true for the amount of processive signals as well as for their velocity (Fig 3B-C). Hence, Upa2 appeared to shuttle on Rrm4-positive transport endosomes [8, 16].

**Figure 3.**
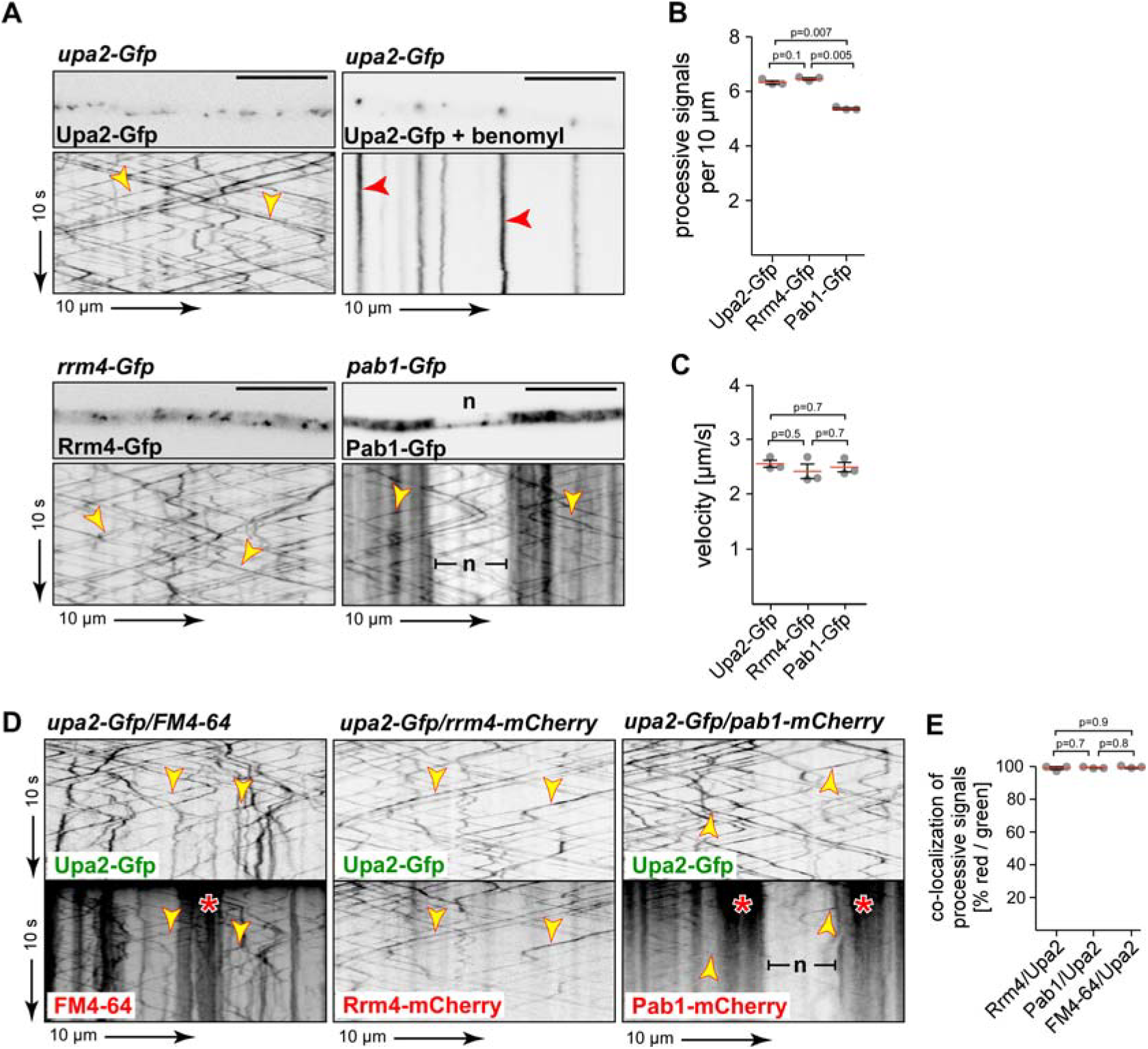
Bidirectional movement of Upa2 on Rrm4-positive endosomes. (**A**) Micrographs (inverted fluorescence image; size bar, 10 µm) and corresponding kymographs of AB33 hyphae (6 h.p.i.) expressing Upa2-Gfp, Rrm4-Gfp or Pab1-Gfp (arrow length on the left and bottom indicate time and distance, respectively). Bidirectional movement is visible as diagonal lines (yellow arrowheads; n, nucleus; Video EV1). After addition of the microtubule inhibitor benomyl static signals are seen as vertical lines (red arrowheads; Video EV1). (**B**) Processive signals per 10 µm of hyphal length (data points representing mean from n = 3 independent experiments, with mean of means, red line, and s.e.m.; unpaired two-tailed Student′s t-test, for each experiment at least 10 hyphae were analysed per strain). (**C**) Velocity of fluorescent signals (velocity of tracks with > 5 µm processive movement; data points representing means from n = 3 independent experiments, with mean of means, red line, and s.e.m.; unpaired two-tailed Student′s t-test; for each experiment at least 30 signals per hypha were analysed out of 10 hyphae per strain). (**D**) Kymographs of AB33 hyphae (6 h.p.i.) expressing pairs of red and green fluorescent proteins as indicated. Fluorescence signals were detected simultaneously using dual-view technology (arrow length as in A). Processive co-localising signals are marked by yellow arrowheads. Areas of static signals are indicated by red asterisks (Video EV2). (**E**) Percentage of processive signals exhibiting co-localisation for strains shown in D. Note, that due to the weaker red fluorescence only clearly detectable red fluorescent signals were analysed for co-localisation with processive green fluorescent signals(data points represent means set to 100% from n = 3 independent experiments, with mean of means, red line, and s.e.m.; unpaired two-tailed Student′s t-test; for each experiment 6 hyphae were analysed per strain).

For experimental verification of endosomal shuttling we stained processive endosomes with the lipophilic dye FM4-64 [10]. Using dual-colour dynamic live imaging, we detected extensive co-localisation of Upa2-Gfp and FM4-64-stained, processively moving endosomes (Fig 3D-E). Furthermore, we tested strains co-expressing Upa2-Gfp and Rrm4-mCherry or Pab1-mCherry (functional C-terminal fusions with the monomeric red fluorescent protein mCherry) [12, 13]. In this instant we also observed extensive co-localisation of Upa2 and Rrm4 (Fig 3D-E, Video EV2) indicating that Upa2 was present on almost all shuttling endosomes [8, 16]. Upa2-Gfp also co-localised with processively moving Pab1-mCherry (Fig 3D-E). However, unlike Rrm4, Pab1-mCherry additionally localised in the cytoplasm resulting in diffuse and static signals that are easily detectable around the region of the nucleus (Fig 3A, D) [12, 13]. Notably, Upa2 did not exhibit this cytoplasmic staining indicating that it is not present in cytoplasmic Pab1-containing complexes, but specifically interacts with endosome-associated Pab1 (Fig 3A, D, E). In essence, Upa2 shuttles exclusively on almost all Rrm4-positive transport endosomes supporting the notion that it is a new component of endosomal mRNPs.

### Endosomal localisation of Upa2 depends on the RNA-binding capacity of Rrm4

Since Upa2 proved to be an endosomal protein, we investigated the functional relationship between Upa2 and Rrm4 by studying the subcellular localisation of endosomal mRNP components in *rrm4*Δ strains. As previously described, the endosomal localisation of Upa1-Gfp is not affected in *rrm4*Δ strains, because its interaction is mRNA-independent and mediated by its FYVE domain (Fig 4A-B, Video EV3) [16]. In contrast, Pab1-Gfp is no longer detectable on shuttling endosomes in *rrm4*Δ strains indicating that without Rrm4 no mRNAs are transported on endosomes (Fig 4A-B, Video EV4) [13]. Interestingly, the endosomal localisation of Upa2 was also no longer detectable in *rrm4*Δ strains (Fig 4A-B, Video EV5) suggesting that the association of Upa2 depends on direct interaction with endosomal mRNP components. In corresponding Western blot experiments, we verified that loss of Rrm4 did not alter the protein amounts of Upa2-Gfp, Pab1-Gfp or Upa1-Gfp (Fig EV2E).

**Figure 4.**
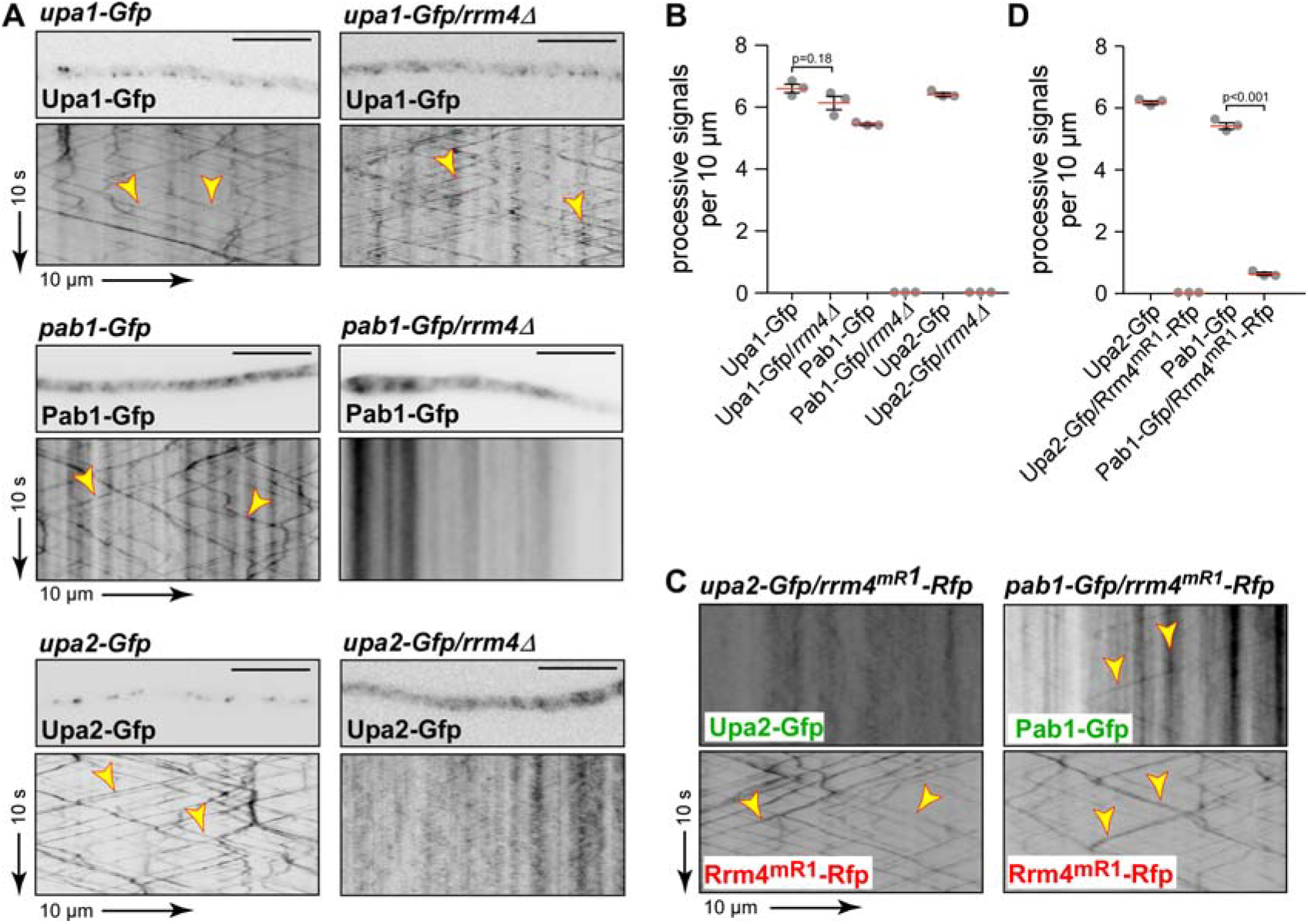
Endosomal localisation of Upa2 is mRNA-dependent. (**A**) Micrographs (inverted fluorescence image; size bar, 10 µm) and corresponding kymographs of AB33 hyphae (6 h.p.i.). Genetic background as indicated above (arrow length on the left and bottom indicate time and distance, respectively). Bidirectional movement is visible as diagonal lines (yellow arrowheads; Videos EV3-EV5). **(B)** Processive signals per 10 µm of hyphal length (data points representing means from n = 3 independent experiments, with mean of means, red line, and s.e.m.; unpaired two-tailed Student′s t-test, for each experiment at least 10 hyphae were analysed per strain). (**C**) Kymographs of AB33 hyphae (6 h.p.i.) expressing pairs of red and green fluorescent proteins as indicated. Fluorescence signals were detected simultaneously using dual-view technology (arrow length as in A). Processive co-localising signals are marked by yellow arrowheads (Videos EV6-EV7). (**D**) Processive signals per 10 µm of hyphal length (data points representing means from n = 3 independent experiments; with mean of means, red line, and s.e.m.; unpaired two-tailed Student′s t-test, for each experiment at least 10 hyphae were analysed per strain).

To investigate a possible mRNA-dependent interaction of Upa2 with shuttling endosomes we studied the effect of the allele Rrm4^mR1^-Rfp (Rrm4 variant with C-terminally fused monomeric Rfp). Previously, it was shown that Rrm4^mR1^ carrying a loss of function mutation in the first RRM domain resulted in drastically reduced RNA binding of Rrm4 [17]. Notably, Upa2 is no longer detectable on endosomes in a Rrm4^mR1^-Rfp background, whereas for Pab1 we observed a strong reduction in movement (Fig 4C-D, Videos EV6-EV7). This indicates that the recruitment of Upa2 to motile transport endosomes is dependent on the RNA-binding capacity of Rrm4. Taken together, although loss of the endosomal components Upa1 and Upa2 results in similar defects during hyphal growth, the mode of endosomal localisation is clearly different. Contrary to Upa1, endosomal localisation of Upa2 is mRNA-dependent, suggesting an interaction with mRNA either directly or indirectly through a protein component of mRNPs such as Pab1 that Upa2 binds via its PAM2 motifs.

### Upa2 carries two functionally important regions, a N-terminal effector domain and a C-terminal GWW motif

Since the mRNA-dependent localisation could be explained by the direct PAM2-dependent interaction with Pab1, we tested the role of the identified PAM2 motifs of Upa2 with regards to function and endosomal localisation. Therefore, we expressed Upa2-Gfp variants in laboratory strain AB33 carrying mutations in one, in multiple and in all identified PAM2 motifs. Surprisingly, none of the PAM2 mutations affected the hyphal growth of *U. maydis* indicating that the interaction of Upa2 with Pab1 is not essential for its role during hyphal growth (Fig 5A-B). Furthermore, endosomal shuttling was not affected by mutating all four PAM2s indicating that the interaction with Pab1 is dispensable for endosomal localisation (Fig EV3A). Hence, Upa2 must contain so far unknown domains critical for endosomal transport functions.

**Figure 5.**
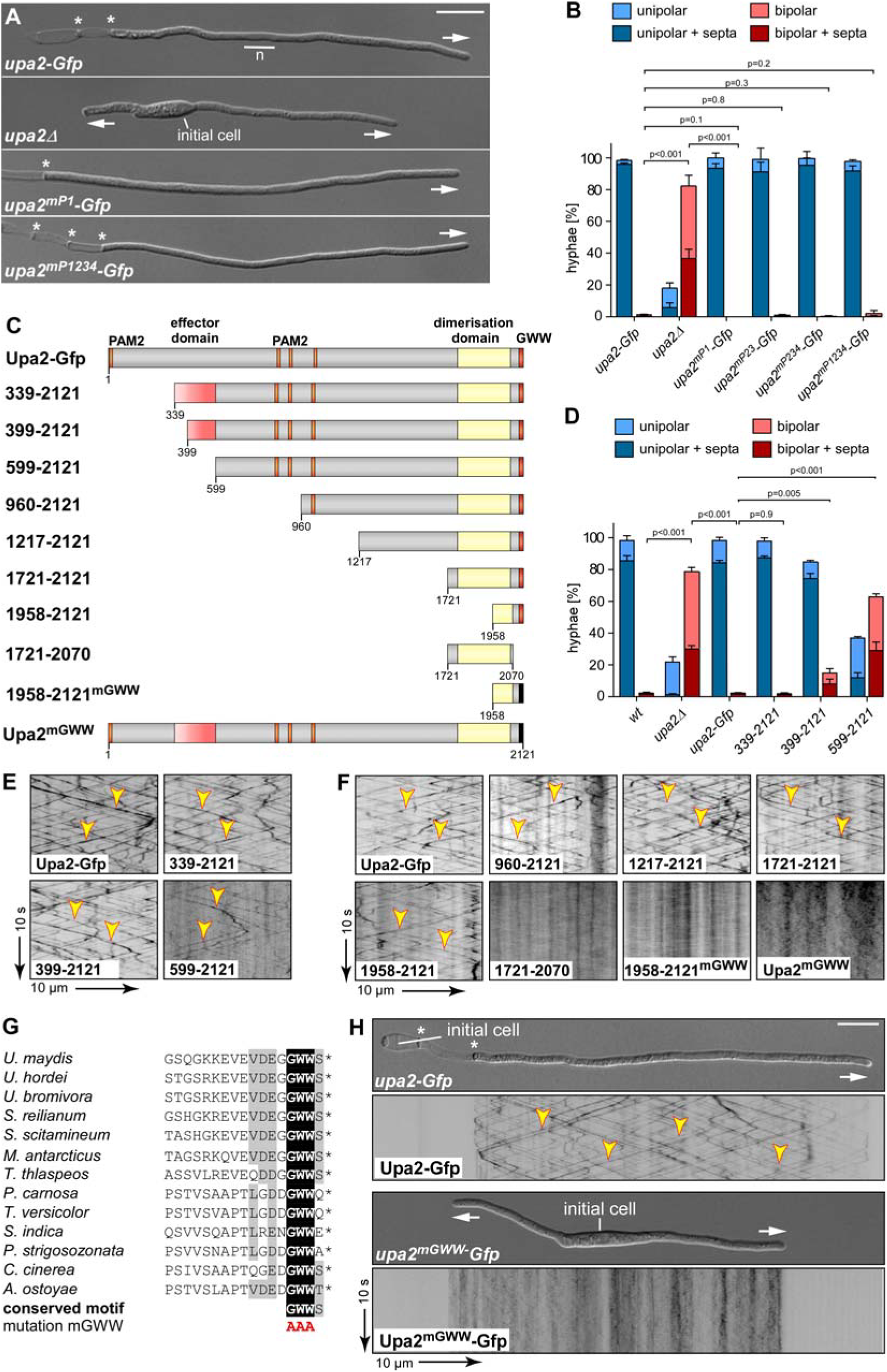
Upa2 carries a functional important effector domain at the N-terminus and a C-terminal GWW motif for endosomal localisation. (**A**) Growth of AB33 derivatives in the hyphal form (6 h.p.i.; growth direction is marked by arrows; asterisks, basal septa; n, nucleus; size bar 10 µm). (**B**) Quantification of hyphal growth (6 h.p.i): unipolarity, bipolarity and septum formation were quantified (error bars, s.e.m.; n = 3 independent experiments, for each experiment >100 hyphae were counted per strain; note that septum formation is given relative to the values of unipolar or bipolar hyphae set to 100%). **(C)** Schematic representation of N-terminal truncations of Upa2 (orange, PAM2 motif; light flow red, effector domain; yellow, dimerization domain; red, GWW; black, mutation in GWW). (**D**) Quantification of hyphal growth of N-terminally truncated Upa2 mutants (6 h.p.i): unipolarity, bipolarity and septum formation were quantified (error bars, s.e.m.; n = 3 independent experiments, for each experiment >100 hyphae were counted per strain; note that septum formation is given relative to the values of unipolar or bipolar hyphae set to 100%). For statistical analysis, the percentage of bipolarity was investigated by using unpaired two-tailed Student′s t-test. Three independent experiments (n = 3) were conducted with at least 100 hyphae per strain. **(E) (F)** Kymographs of AB33 hyphae (6 h.p.i.); genetic background as indicated (arrow length on the left and bottom indicate time and distance, respectively). Bidirectional movement is visible as diagonal lines (yellow arrowheads). **(G)** Comparison of the C-terminal amino acid sequence of Upa2 in *U. maydis* and related fungal species. Accession numbers can be found in Appendix Table S7. **(H)** Growth of AB33 derivatives in the hyphal form (6 h.p.i.; growth direction is marked by arrows; asterisks, basal septa; size bar, 10 µm) and corresponding kymographs of AB33 hyphae (6 h.p.i.). Genetic background as indicated in left bottom panel (arrow length on the left and bottom indicate time and distance, respectively). Bidirectional movement is visible as diagonal lines (yellow arrowheads; Video EV8).

To map these domains, we studied N-terminal truncations carrying C-terminal Gfp fusions (Fig 5C). Respective alleles were integrated at the homologous locus replacing the endogenous copy of *upa2*. Expression was driven by the native *upa2* promoter. Removal of the first 338 amino acids did not interfere with the function of the protein (Upa2-339-2121; Fig 5C-D). However, additional truncations resulted in increase of bipolar hyphae indicating reduced (Upa2-399-2121) or loss of function (Upa2-599-2121, 960-2121, 1217-2121, 1721-2121 and 1958-2121; Fig 5C-D, Fig EV3B). Importantly, protein amounts (e.g. 399-2121 and 399-2121; Fig EV3C) and endosomal shuttling of these versions were not drastically affected (Fig 5E-F; Fig EV3D-E). Hence, a currently unknown effector domain important for endosomal mRNA transport is located between amino acid position 339 and 599. Notably, the non-functional variant Upa2-599-2121 still contained three PAM2 motifs, consistent with the finding that the interaction with Pab1 is not sufficient for function.

For mapping of the domain that mediates endosomal localisation of Upa2, we assayed additional N-terminal truncations. All variants still shuttled indicating that the last 163 amino acids were sufficient for endosomal shuttling (Fig 5C, E-F). Upa2-1217-2121 did no longer contain any PAM2 region for interaction with Pab1 (Fig 5C). This was consistent with the finding that the PAM2 motifs of Upa2 were not essential for endosomal localisation. Closer inspection by sequence comparison revealed a conserved GWW sequence at the very C-terminus (Fig 5G), which was shown in other proteins to function in protein/protein interaction [21, 22]. Mutating this short sequence resulted in loss of shuttling without drastic changes in protein amounts (Fig 5F, Fig EV3F). This holds true when testing the mutation in the context of the full length protein (Fig 5H, Video EV8). Importantly, we also observed a loss of function phenotype for this mutation, i.e. an increased number of bipolar hyphae (Fig 5H, Fig EV3G). This was not due to altered protein amounts (Fig EV3F). Thus, the conserved C-terminal GWW motif is essential for endosomal localisation of Upa2 and endosomal localisation is important for the function of the protein. In essence, besides the PAM2 motifs, Upa2 contains a functionally important N-terminal effector domain and a C-terminal GWW motif for interaction with an endosomal mRNP component (see Discussion).

### Loss of Upa2 causes defects in the formation of endosomal mRNPs

Finally, we studied the role of Upa2 during endosomal transport in closer detail. Observation of endosomal shuttling of Rrm4-Gfp, Upa1-Gfp and Rab5a-Gfp revealed that loss of Upa2 did not cause drastic alterations in bidirectional movement. We did notice a slight increase in the velocity and amount of shuttling signals (Fig 6A-C, Videos EV9-EV12). In case of Rrm4-Gfp we observed that a substantial fraction of the protein also stained structures that resembled microtubule bundles (Fig 6A; Fig EV4B). Consistently, in benomyl treated hyphae bundle-like Rrm4-Gfp signals were no longer detected (Fig EV4B-C). This suggests that Rrm4 localisation is disturbed in the absence of Upa2.

**Figure 6.**
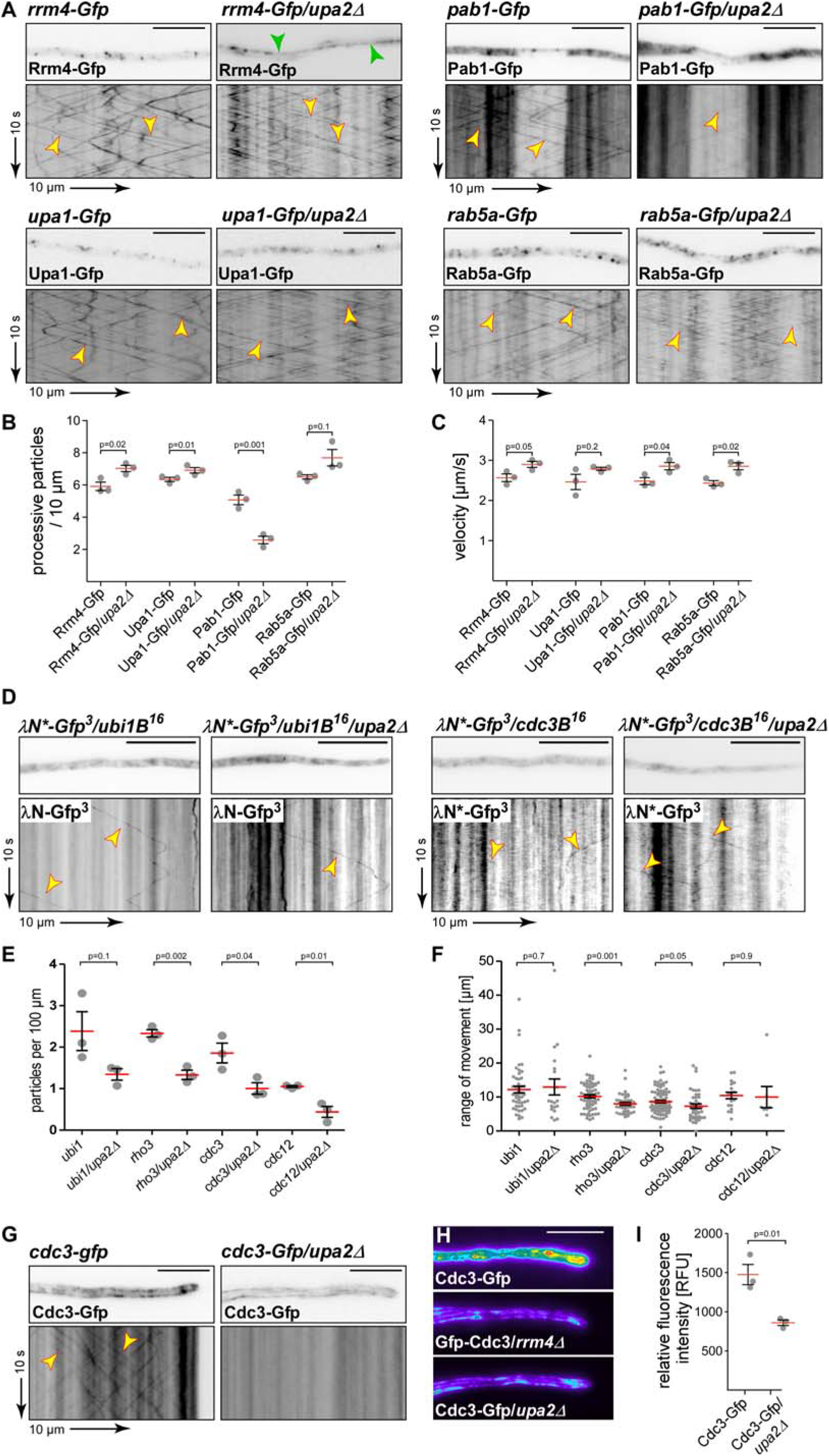
Loss of Upa2 causes defects in the formation of endosomal mRNPs. **(A)** Micrographs of hyphal tips (inverted fluorescence image; size bar, 10 µm) and corresponding kymographs of AB33 hyphae (6 h.p.i.). Genetic background as indicated above (arrow length on the left and bottom indicate time and distance, respectively). Bidirectional movement is visible as diagonal lines (yellow arrowheads; size bar, 10 µm; Videos EV9-12). **(B)** Processive signals per 10 µm of hyphal length (data points representing means from n = 3 independent experiments, with mean of means, red line, and s.e.m.; unpaired two-tailed Student′s t-test. For each experiment at least 10 hyphae per strain were analysed. (**C**) Velocity of fluorescent signals (velocity of tracks with > 5 µm processive movement; data points representing means from n = 3 independent experiments; with mean of means, red line, and s.e.m.; unpaired two-tailed Student′s t-test, for each experiment at least 30 signals per hypha were analysed out of 10 hyphae per strain). (**D**) Micrographs of hyphae and corresponding kymographs of AB33 derivatives (8 h.p.i.). Genetic background as indicated above (arrow length on the left and bottom indicate time and distance, respectively). Bidirectional movement of mRNA is visible as diagonal lines (yellow arrowheads; size bar, 10 µm; Video EV13). (**E**) Number of particles per 100 µm hypha (data points representing means from n = 3 independent experiments, with mean of means, red line, and s.e.m.; unpaired two-tailed Student′s t-test, for each experiment at least 7 hyphae were analysed per strain). (**F**) Range of movement of motile mRNAs (red line, median; n = 3 independent experiments; unpaired Student′s t-test). (**G**) Micrographs of hyphal tips (inverted fluorescence image; size bar, 10 µm) and corresponding kymographs of AB33 hyphae (6 h.p.i.). Genetic background as indicated above (arrow length on the left and bottom indicate time and distance, respectively). Bidirectional movement is visible as diagonal lines (yellow arrowheads; size bar, 10µm; Video EV14). (**H**) Heat maps of hyphal tips (6 h.p.i.) of AB33 derivatives expressing Cdc3-Gfp comparing wild type (top panel) with *rrm4Δ* and *upa2Δ* strains (middle, bottom panel), indicating relative fluorescence intensity differences (maximum projection of z-stacks; size bar, 10 µm; red/yellow to green/blue, high to low intensities). (**I**) Relative fluorescence intensity (RFU), first 10 µm from hyphal tip (data points representing means from n = 3 independent experiments, with mean of means, red line, and s.e.m.; unpaired two-tailed Student′s t-test, for each experiment at least 10 hyphae were analysed per strain).

Interestingly, studying shuttling of Pab1-Gfp revealed that the amount of Pab1-positive endosomes was reduced to about 50% (Fig. 6A-B), suggesting that Upa2 is needed for an efficient interaction of Pab1-containing mRNAs with endosomes. To address this notion more directly we applied RNA live imaging [23]. We studied four different target mRNAs of Rrm4-dependent endosomal mRNA transport encoding the ubiquitin fusion protein Ubi1, the small G protein Rho3, as well as the septins Cdc3 and Cdc12 (Fig EV4D) [12–14]. Importantly, in the absence of Upa2 we observed in all four cases that the number of processively transported mRNAs was reduced to about 50% (Fig 6D-E). This is consistent with the shuttling of Pab1-Gfp that was reduced to comparable extend (Fig 6A-B). The range of movement (Fig 6F) and the velocity (Fig EV4E) of transported *ubi1*, *rho3*, *cdc3* and *cdc12* mRNA particles was not changed significantly. Thus, loss of Upa2 affects most likely assembly or long-term association of mRNPs for transport.

An important function of endosomal septin mRNA transport is the local assembly of septin heteromeric subunits that are transported to hyphal growth poles to form higher-order filaments with a gradient emanating from this pole [12, 14]. To address the role of Upa2 we used a functional septin fusion protein Cdc3-Gfp as read-out (Cdc3 carrying eGfp at its N-terminus; notably the 5′ and 3′ untranslated regions of *cdc3* were preserved to keep potential regulatory elements and Rrm4 binding sites intact) [12, 14]. Loss of Upa2 abolished endosomal shuttling of Cdc3-Gfp and as an expected consequence also the Cdc3-Gfp containing gradient at hyphal tips was no longer detectable (Fig 6G-I, Video EV14). The strong reduction of septin protein on the surface of endosomes is consistent with the fact that stable endosomal localisation of septin subunits depend on each other[14]. Since *cdc3* and *cdc12* mRNAs are both reduced to 50% the amount of translated protein appears to be insufficient to support endosomal localisation (see Discussion). In essence, Upa2 is essential for the correct association of mRNAs, Pab1, as well as Rrm4, on endosomes. Thus, we identified an endosome-specific and functionally important factor that functions as novel core component of endosomal mRNA transport.

## Discussion

Functional units of mRNA transport are higher-order mRNPs consisting of cargo mRNAs, RNA-binding proteins and additional interacting proteins [5]. A crucial task is to differentiate between core components that are essential to orchestrate transport, and accessory components. The latter might mediate for example translational regulation of cargo mRNAs. Here, we identified Upa2 as a novel core component of endosomal mRNA transport. This interactor of the poly(A)-binding protein appears to be important for assembly or stabilisation of higher-order mRNPs for endosomal transport suggesting that it functions as a scaffold protein (Fig 7).

**Figure 7.**
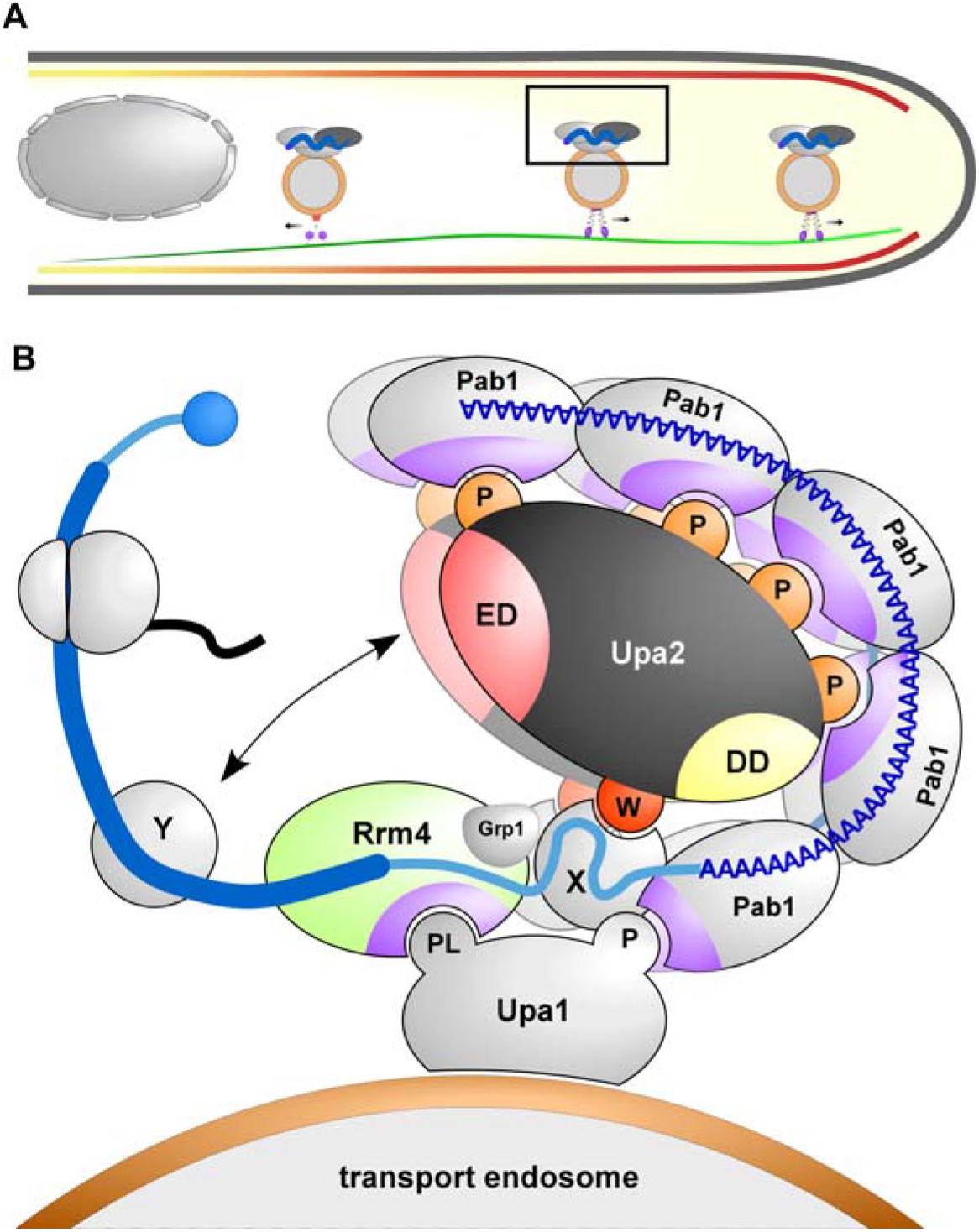
Model depicting Upa2 as novel core component of endosomal mRNP transport. (**A**) Schematic drawing of infectious hypha (red, septin filaments with gradient; green, microtubules; grey circles with golden edge, transport endosomes with mRNPs and molecular motors attached; boxed region is enlarged in B). (**B**) Simplified view of known mRNP components at the cytoplasmic surface of transport endosome and a proposed interplay. Upa2 harbours at least four different protein/protein interfaces: <u>P</u>AM2 motifs (orange), <u>e</u>ffector <u>d</u>omain (light red), <u>d</u>imerisation <u>d</u>omain (yellow), G<u>W</u>W motif (dark red). MLLE domains of Pab1 and Rrm4 are given in purple. Additional symbols: 5′ cap, blue circle; mRNA as blue line with ORF (thick line); translating ribosome (grey) with nascent chain (black). Additional mRNP components that are predicted to interact with Upa2 are given as grey circles labelled X and Y (for further details, see text).

### A novel multi PAM2 protein for endosomal mRNA transport

Upa2 is a large protein of 232 kDa that contains four functional PAM2 motifs for interaction with the C-terminal MLLE domain of Pab1. It is known from earlier structural studies that the C-terminal MLLE domain of human PABC1 forms a defined peptide pocket to accommodate various PAM2 sequences [24]. Currently, only three MLLE-containing proteins are described: the poly(A)-binding protein, the E3 ubiquitin ligase UBR5 and mRNA transporter Rrm4. UBR5 is a HECT-type (<u>h</u>omologous to the <u>E</u>6AP <u>C</u> terminus) ligase that functions in translational regulation and microRNA-mediated gene silencing [25].

The PAM2 motif is present in various PABC1-interaction partners such as eRF3, GW182 and PAN3 functioning in various steps of posttranscriptional control such as translation, miRISC assembly and deadenylation, respectively [26]. The vast majority of proteins harbour a single PAM2 motif, only eRF3 and Tob contain two overlapping PAM2 motifs for MLLE interaction [27, 28]. Therefore, it is exceptional to find four PAM2 motifs in Upa2 and due to the potential dimerisation via its C-terminal coiled coil region Upa2 offers an extensive interaction platform for Pab1 (Fig 7).

Experimental evidence confirmed that the PAM2 motifs of Upa2 interact with Pab1. However, it seems to be an utter paradox that these evolutionarily conserved PAM2 motifs are not important for the function of Upa2 during endosomal mRNA transport. The same holds true for Upa1, where the PAM2 motif is also dispensable for function [16]. A possible explanation is a high level of redundancy. This is supported by the fact that in endosomal mRNPs there are verified PAM2 motifs in Upa1 and Upa2 as well as additional cryptic versions in Rrm4. It is conceivable that all PAM2 sequences and their variants must be mutated to observe defects in mRNA transport and consequently in hyphal growth. In fact, redundancy was already observed during the study of PAM2-like motifs. Only deletion of both PAM2-like sequences in Upa1 resulted in loss of function [16]. Alternatively, we might not be able to detect the functional significance of the PAM2 / Pab1 interaction under optimal laboratory conditions. The interaction could be of specific importance under certain stress conditions. Comparably, a function of the small glycine rich protein Grp1 was only observed during hyphal growth under cell wall stress conditions [15].

Besides PAM2 motifs and a coiled coil region for dimerization, we identified two additional functionally important regions. Firstly, we succeeded in mapping a novel N-terminal effector domain. Interestingly, this domain is embedded in a part of the protein that is predicted to be rich in intrinsically disordered regions (IDRs; Fig EV5). In the future, we need to carry out an extensive fine mapping to identify clear boundaries of this domain and identify the respective interaction partner. Secondly, we discovered a C-terminal GWW motif that is essential for the endosomal localisation of Upa2. Since the presence of Upa2 depends on the RNA-binding capacity of Rrm4, we assume that the GWW motif does not bind endosomal lipids directly. Most likely, it interacts with a protein component of endosomal mRNPs (Fig 7). This assumption is supported by the fact that an evolutionarily conserved GWW motif at the C-terminus of bacterial endoribonuclease RNase E interacts with a peptide-binding pocket of a PNPase (exoribonuclease polynucleotide phosphorylase; 21, 22]. Furthermore, GWW motifs have been implicated in intramolecular interactions of two SH3 domains of the NADPH oxidase component p47^phox^ [29]. Related dispersed GW motifs in the posttranslational regulator GW182 mediate recruitment of CCR4-NOT deadenylation components during miRNA-mediated repression [30]. In essence, we identified at least four different types of interfaces for protein/protein interactions in Upa2 supporting the hypothesis that it serves as a scaffold protein.

### The function of Upa2 during endosomal transport

In order to study the function of Upa2 *in vivo* we combined genetic with cell biological approaches. Loss of Upa2 causes the same aberrant growth phenotype and reduced Cts1 secretion as observed in *rrm4*Δ and *upa1*Δ strains [13, 16], suggesting that these proteins function in the same pathway. This is supported by a phylogenetic analysis showing the co-appearance of all three proteins.

The bipolar hyphal growth phenotype can in part be explained by the defects in forming the gradient of septin filaments at the growth pole, since septins are important during the initial phase of unipolar hyphal growth [12, 14]. The formation of the septin gradient depends on the endosomal transport of septin heteromers and in turn heteromer assembly depends on endosomal transport of the septin encoding mRNAs [12, 14]. Thus, the most rational explanation for the observed defects in hyphal growth in *upa2*Δ strains is the reduced transport of septin mRNAs resulting in fewer septin proteins on endosomes. Since the presence of the different septins on transport endosomes are interdependent [14], the reduced amount might not be able to support endosomal assembly and localisation of septin proteins causing the observed defects in septin gradients. The hypothesis of local translation on the endosomal surface for septin assembly [12, 14] has recently been supported by the finding that most heteromeric protein complexes are co-translationally assembled in eukaryotes [31].

Loss of Upa2 also causes a slight increase in the amount and velocity of transport endosomes suggesting that the absence of Upa2 and the reduced associated mRNA cargos has an influence on transport endosomes in general. However, the most profound differences were recognized studying the mRNP components Rrm4 and Pab1. We observed an aberrantly strong formation of Rrm4 on microtubules and reduced endosomal localisation of Pab1. Hence, Upa2 appears to be crucial for the stable association of mRNPs on the surface of endosomes during transport. This is consistent with important key findings of this study that Upa2, like Rrm4, specifically localises to transport endosomes and that it is present an almost all transport endosomes. In conjunction with the aforementioned scaffolding function, Upa2 fulfils the necessary criteria to function as a novel core component of endosomal mRNP transport.

### Conclusions

In recent years the identification of components of the endosomal mRNA transport machinery in *U. maydis* has advanced significantly (Fig 7). Importantly, it is the first system, where a transcriptome-wide binding landscape of the key RBP is available. Comparing bound RNAs of Rrm4 with the accessory component Grp1 revealed that Rrm4 orchestrates a tailored transport strategy for distinct sets of cargo mRNAs [15]. Here, we add the evolutionarily conserved Upa2 as an important novel piece to our jigsaw puzzle of endosomal mRNA transport.

Key concepts appear to be conserved throughout evolution and might also be applicable to higher eukaryotes. In fungi, for example, Rrm4, Upa1 and Upa2 are conserved throughout Basidiomycetes suggesting that endosomal mRNA transport is more wide-spread than currently anticipated [20]. Consistently, in animal systems, endosomal components have been implicated in mRNA transport during axonal growth [32, 33] and mRNAs associated with Rab5-positive endosomes were described in plants [34]. More recently, endosomal mRNA transport and coupled translation at late endosomes were described to be crucial for mitochondrial function in polar growing axons [35]. Hence, endosome-coupled translation seems to be conserved from fungi to men and is now critical to obtain insights in the mechanisms of how mRNPs stably associate with endosomes [36]. Finally, discovering the key RNA-binding protein Rrm4 that is linked intensively and intimately with endosomal membranes fits to the new emerging concept that RNA and membrane biology are tightly intertwined. Membrane-associated RBPs (memRBPs) are most likely important at all intracellular membranes to orchestrate spatio-temporal expression [37]. These findings stress the vital role of *U. maydis* as a model for RNA biology [4, 10].

## Materials and methods

### Plasmids, strains and growth conditions

For cloning of plasmids and GST pulldown experiments, *E. coli* Top10 cells (Life Technologies, Carlsbad, CA, USA) and *E. coli* Rosetta2 pLysS (Merck 71403) were used, respectively. Transformation, cultivation and plasmid isolation were conducted using standard techniques. All *U. maydis* strains are derivatives of AB33, in which hyphal growth can be induced [18]. Yeast-like cells were incubated in complete medium (CM) supplemented with 1% glucose, whereas hyphal growth was induced by changing to nitrate minimal medium (NM) supplemented with 1% glucose, both at 28°C [18]. Detailed growth conditions and cloning strategies for *U. maydis* are described elsewhere [8, 38, 39]. All plasmids were verified by sequencing. Strains were generated by transforming progenitor strains with linearised plasmids. Successful integration of constructs was verified by diagnostic PCR and by Southern blot analysis [38]. For ectopic integration, plasmids were linearised with SspI and targeted to the *ip*^*S*^ locus [40]. Wild-type strain UM521 genomic DNA was used as a template for PCR amplifications of ORFs, unless otherwise stated. Yeast two-hybrid tests were carried out using *S. cerevisiae* strain AH109 (Clontech Laboratories Inc., Mountain View, CA, USA). A detailed description of all plasmids and strains is given in Appendix Tables S1 to S6. Sequences are available upon request.

### Microscopy, image processing and image analysis

Laser-based epifluorescence-microscopy was performed on a Zeiss Axio Observer.Z1 equipped with CoolSNAP HQ2 CCD (Photometrics, Tuscon, AZ, USA) and ORCA-Flash4.0 V2+ CMOS (Hamamatsu Photonics Deutschland GmbH, Geldern, Germany) cameras. For excitation we used a VS-LMS4 Laser-Merge-System (Visitron Systems, Puchheim, Germany) that combines solid state lasers for excitation of Gfp (488 nm at 50 or 100 mW) and Rfp/mCherry (561 nm at 50 or 150 mW).

For the quantification of unipolar hyphal growth, cells were grown in 20 ml cultures to an OD_600_ of 0.5 and hyphal growth was induced. After 6 hours, more than 100 cells per experiment were imaged and analysed for growth behaviour. Cells were scored for unipolar and bipolar growth as well for formation of empty sections. At least three independent experiments were conducted. For statistical analysis, the rate of bipolarity was investigated by using unpaired two-tailed Student′s t-test.

For analysis of signal number and velocity, we recorded videos with an exposure time of 150 ms and 150 frames taken. All videos and images were processed and analysed using Metamorph (Version 7.7.0.0, Molecular Devices, Seattle, IL, USA). Kymographs were generated using a built-in plugin and processively moving particles were counted manually. The average velocity was determined by quantifying processive signals (movement > 5 µm). Data points represent means from three independent experiments (n = 3) with mean of means (red line) and s.e.m. For each experiment at least 30 signals per hypha were analysed out of 10 hyphae per strain.

For quantification of Cdc3 fluorescence intensity line-scans were conducted in a region of 10 µm from hyphal tips. Relative fluorescence intensities of 30 hyphal tips per strain were averaged. Data points represent means from three independent experiments (n = 3) with mean of means (red line) and s.e.m.. Colocalization studies of dynamic processes were carried out by using a two-channel imaging system (DV2, Photometrics, Tucson, AZ, USA) [16, 41].

### RNA live imaging, FM4-64 staining and benomyl treatment

RNA imaging in living cells was conducted by using the λN-based green-RNA method described previously [12, 14]. For RNA visualization of *ubi1* and *rho3* λN was fused to three copies of enhanced Gfp and 16 copies of the boxB loop were inserted in the 3′ UTR of the respective mRNAs. Expression was driven either by the native *ubi1* promoter or in case of *rho3* by the constitutively active promoter P_otef_. For RNA visualization of *cdc3* and *cdc12* a modified λN (λN*) was fused to three copies of enhanced Gfp and 16 copies of the boxB loop were inserted in the 3′ UTR of the respective mRNAs. Both constructs were under the control of the constitutive active promoter P_otef_. Three independent experiments (n = 3) were conducted with at least 10 hyphae per strain. Statistical tests were performed using Graphpad Prism5 (version 5.00; Graphpad Software, La Jolla, CA, USA). A detailed protocol for subsequent data analysis was described elsewhere [14]. For staining of cells with FM4-64, a 1 ml sample of hyphal cells was labelled with 0.8 µM FM4-64 (Thermo Fisher, Waltham, MA, USA). After incubation for 1 min at room temperature, the labelled cells were analysed by fluorescence microscopy. For benomyl treatment, a 1 ml sample of hyphal cells was treated with 20 µM benomyl (Sigma-Aldrich, Taufkirchen, Germany). After incubation for 30 min at room temperature with agitation samples were analysed by microscopy.

### Fluorimetric measurement of chitinolytic activity

Chitinolytic activity measurements of *U. maydis* cells were carried out as described elsewhere [16, 19]. Briefly, *U. maydis* cell suspensions were grown to an OD_600_ of 0.5. The culture was divided in half, yeast-like growing cells were measured directly while activity of hyphae was measured 6 h after induction of hyphal growth. 30 µl of the culture were mixed with 70 µl 0.25 µM 4-Methylumbelliferyl-β-D-N,N‘,N‘‘-triacetylchitotrioside (MUC, Sigma-Aldrich, Taufkirchen, Germany), a fluorogenic substrate for chitinolytic activity. After incubation for 1 h, the reaction was stopped by addition of 200 µl 1M Na_2_CO_3_, followed by detection of the fluorescent product with the fluorescence spectrometer Infinite M200 (Tecan Group Ltd., Männedorf, Switzerland) using an excitation and emission wavelength of 360 nm and 450 nm, respectively. Chitinase activity was set and reported in relation to AB33 (wt) activity. Five independent biological experiments were performed with three technical replicates per strain.

### Yeast two-hybrid analysis

The yeast two-hybrid analyses were carried out as described elsewhere [16]. Briefly, using the two-hybrid system Matchmaker 3 from Clontech, strain AH109 was co-transformed with derivates of pGBKT7-DS and pGADT7-Sfi (Appendix Table S4) and cells were grown on synthetic dropout (SD) plates without leucine and tryptophan at 28° C for 4 days. Subsequently, colonies were patched on SD plates without leucine and tryptophan (control) or on SD plates without leucine, tryptophan, histidine and adenine (selection). Plates were incubated at 28°C for 3 days to test for growth under selection condition. For qualitative plate assays cells were cultivated in SD without leucine and tryptophan to OD_600_ of 0.5 and successively diluted with sterile water in four steps at 1:5 each. 4 µl were spotted on control as well as selection plates and incubated at 28° C for 3 days. Colony growth was documented with a LAS 4000 imaging system (GE Healthcare Life Sciences, Little Chalfont, UK).

### Protein extracts and Western blot analysis

*U. maydis* hyphae were harvested 6 hours post induction (h.p.i.) by centrifugation (7546 × g, 10 minutes) and resuspended in 2 ml of either urea buffer (8 M Urea, 50 mM Tris/HCl pH8; Fig EV2E and Fig EV3F, right panel) or l-arginine rich buffer (0.4 M sorbitol; 5 % glycerol; 50 mM Tris/HCl pH7.4; 300 mM NaCl; 1 mM EDTA; 0.5% Nonidet P-40; 0.1% SDS; 72.5 mM l-arginine, Fig EV3C and Fig EV3F, left panel) supplemented with protease inhibitors (1 tablet of complete protease inhibitor per 25 ml, Roche, Mannheim, Germany; 1 mM DTT; 0.1 M PMSF; 0.5 M benzamidine). Cells were lysed in a Retsch ball mill (MM400; Retsch, Haan, Germany) while keeping samples constantly frozen using liquid nitrogen. 2 ml cell suspension per grinding jar with two grinding balls (d = 12 mm) were agitated for 10 minutes at 30 Hz. Protein concentrations were measured by Bradford assay (Bio-Rad, Munich, Germany) and samples were adjusted to equal amounts. For Western Blot analysis, protein samples were supplemented with Laemmli buffer and heated to 60°C-80°C for 10 minutes, resolved by 8% SDS-PAGE and transferred to a nitrocellulose membrane (GE Healthcare, Munich, Germany) by semi-dry blotting. Membranes were probed with α-Gfp (Roche, Freiburg, Germany) and α-actin (MP Biomedicals, Eschwege, Germany) antibodies. As secondary antibody a mouse IgG HRP conjugate was used (Promega, Madison, WI, USA). Detection was carried out by using AceGlow (VWR Peqlab, Erlangen, Germany).

### GST pull down experiments

Derivatives of plasmids pGEX and pET15B (Appendix Table S5) were transformed into *E. coli* Rosetta2 pLysS (Merck 71403). Overnight cultures were diluted 1:50 in a final volume of 50 ml. Protein expression was induced with 1mM IPTG for 4 h at 28°C. Cells were pelleted, resuspended in 10 ml lysis buffer and lysed by sonication. Cell lysate was centrifuged at 16,000 g for 15 minutes and the supernatant was transferred to the fresh microcentrifuge tubes. 50 µl glutathione beads (GE Healthcare) were transferred to a new 1.5 ml microcentrifuge tube and washed 3 times with 1 ml lysis buffer (20 mM Tris-Cl, pH 7.5; 200 mM NaCl; 1mM EDTA, pH 8.0; 0.5 % Nonidet P-40; 1 tablet complete protease inhibitor per 50ml, Roche, Mannheim, Germany). For each pulldown, 1 ml of supernatant of GST-tagged protein was added to the washed beads, incubated for 1 h at 4° C and subsequently washed 5 times with 1 ml lysis buffer. 1 ml supernatant containing His-tagged Upa2 variant was added directly to the washed GST bead and incubated for 1 h at 4° C and subsequently washed 5 times with lysis buffer. The beads were boiled for 6 min at 99° C in 100 µl of 1× Laemmli buffer. 10 µl of each GST-pulldown fraction was analysed by SDS-PAGE and Coomassie blue (CBB R250) staining. 1 ml supernatant containing Upa2 variant was added directly to the washed Ni-NTA (Macherey-Nagel) bead, incubated for 1 h at 4° C and subsequently washed 5 times with 1 ml lysis buffer. 20 µl of each Upa2 variants were loaded on control lanes in SDS PAGE as an input. Note, that the input fraction “H” shows Ni-NTA precipitated Upa2, which could not be visualized before enrichment.

For Western blotting protein samples were resolved by 10% SDS-PAGE and transferred to a PVDF membrane (GE Healthcare) by semi-dry blotting. Western blot analysis was performed with α-GST (Sigma G7781), α-His (Sigma H1029). α-Rabbit IgG HRP conjugate (Promega W4011), α-mouse IgG HRP conjugate (Promega W4021) were used as secondary antibodies. Activity was detected using the AceGlow blotting detection system (VWR Peqlab, Erlangen, Germany).

### Phylogenetic analysis and bioinformatics

Sequence data for *U. maydis* genes was retrieved from the PEDANT database (http://pedant.gsf.de/). Accession numbers of *U. maydis* genes used in this study: *upa2* (UMAG_10350), *rrm4* (UMAG_10836), *pab1* (UMAG_03494), *upa1* (UMAG_12183), *rab5a* (UMAG_10615) and *cdc3* (UMAG_10503). Orthologs were identified using fungiDB [42, 43], Ensembl Fungi [44] and NCBI blastp tool [45]. Sequences alignments were performed with ClustalX (version 2.0.12) [46]. Domains were predicted using SMART [47, 48], conserved domain database from the NCBI (CDD) [49] and active search using ScanProsite [50]. The coiled coil dimerisation domain was predicted using the Interpro COILS program [51]. The phylogenetic tree is based on the NCBI taxonomy and was created using phyloT online tool (phylot.biobyte.de). The lengths of the lines do not represent evolutionary distance. The accession numbers for the proteins can be found in Appendix Table S7).

Intrinsically disordered regions were predicted using the PONDR VL3-BA algorithm (www.pondr.com, Molecular Kinetics, Inc., IN, USA). VL3-BA is a feedforward neural network predictor generating outputs between 0 and 1 which are smoothed over a sliding window of 9 amino acids. Regions with values of 0.5 or above are considered disordered and are marked in red, while peptide regions with values lower than 0.5 are considered ordered and are marked in blue.

## Acknowledgements

We acknowledge Drs. A. Brachmann, J. Béthune, K. Zarnack and lab members for discussion and reading of the manuscript. We are grateful to U. Gengenbacher for excellent technical assistance. The work was funded by grants from the Deutsche Forschungsgemeinschaft to MF (FE448/5-2 DFG-FOR1334; FE448/10-1 DFG-FOR2333; DFG-CRC1208 project number 267205415 and CEPLAS EXC1028).

## Author contributions

SJ, TP, SB and MF designed this study and analysed the data. SJ and TP contributed equally to the genetic and cell biological analysis of Upa2. SB carried out the initial cell biological characterisation of Upa2. KM and SZ performed RNA live imaging experiments. SKD analysed the PAM2/MLLE interaction *in vitro*. MF, SJ and TP drafted and revised the manuscript with input from all co-authors. MF and TP directed the project.

---

**Figure EV1.**
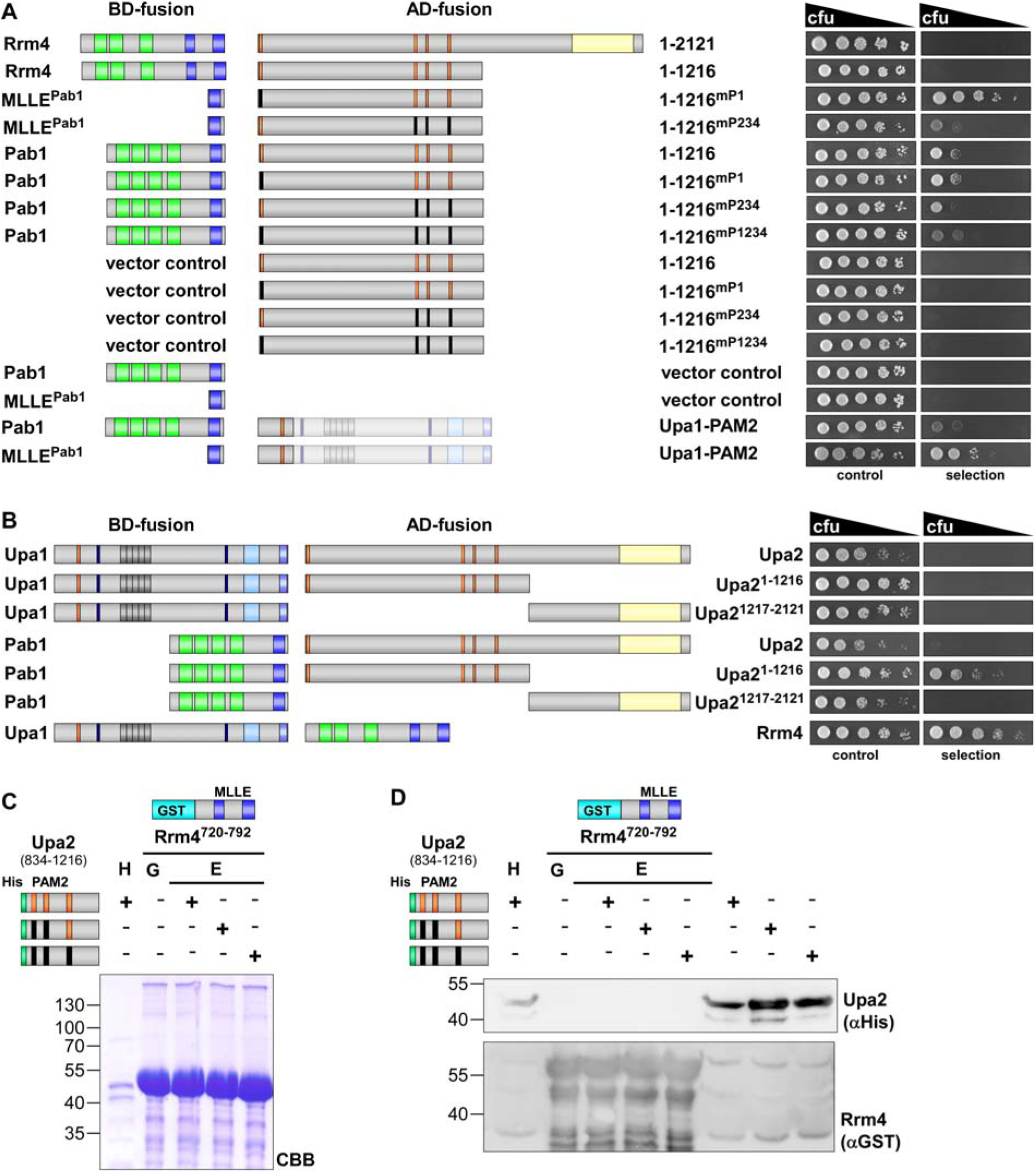
PAM2 motifs of Upa2 do not interact with MLLE domains of Rrm4. (**A-B**) Two-hybrid analysis with schematic representation of protein variants tested on the left (Colour scheme as in Fig 1A, C). Yeast cultures were serially diluted 1:5 (decreasing colony forming units, cfu) and spotted on respective growth plates controlling transformation and assaying reporter gene expression (see Materials and methods). (**C**) GST co-purification experiments with components expressed in *E. coli*: N-terminal His_6_-tagged versions of Upa2^834-1216^, Upa2^834-1216_mP23^, and Upa2^834-1216_mP234^ (amino acids 834–1216; mutation in the PAM2 motifs number 2 and 3 as well as 2, 3 and 4, respectively) were tested for binding to the MLLE domain of Rrm4 (GST fusion to Rrm4^720-792^ containing two MLLE domains). Lane “H” shows Ni-NTA precipitated His_6_-Upa2, lane “G” shows input of GST-Rrm4720-792). After GST affinity chromatography proteins were eluted (lanes marked with “E”). Interaction studies were performed with the soluble fraction of protein extracts from *E. coli* to demonstrate specific binding. (**D**) GST co-purification experiments as shown in B. The absence of Upa2^834-1216^ interaction with the MLLE domains of Rrm4 was demonstrated in more sensitive Western blot experiments (used antibodies are given on the right). The three lanes on the right show all Upa2 versions after Ni-NTA precipitation.

**Figure EV2.**
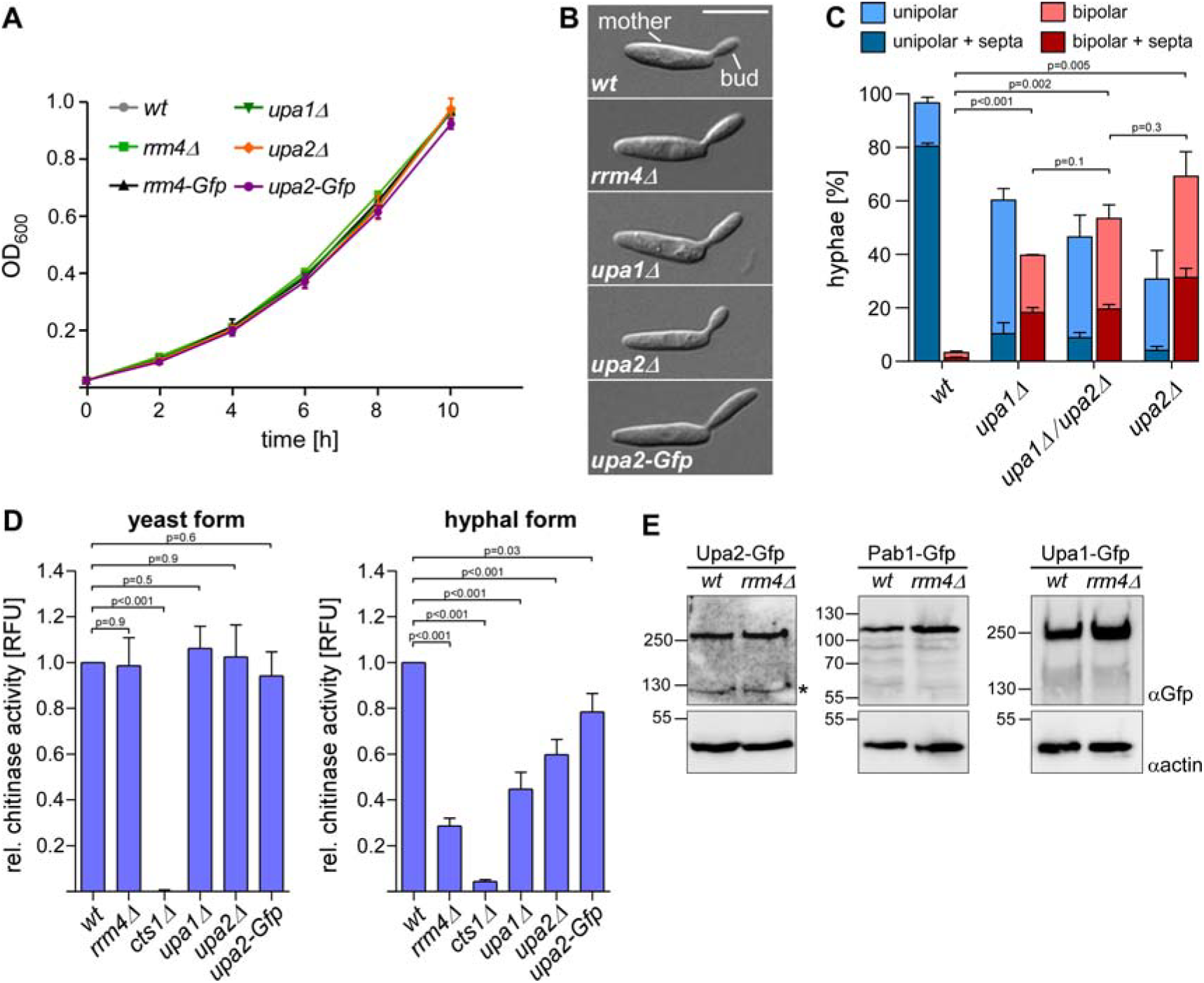
Loss of Upa2 causes defects in secretion of chitinase Cts1. (**A**) Growth of yeast cells of indicated strains in liquid culture. (**B**) DIC images of yeast cells (size bar, 10 µm). (**C**) Quantification of hyphal growth (6 h.p.i): unipolarity, bipolarity and septum formation were quantified (error bars, s.e.m.; n = 3 independent experiments, for each experiment >100 hyphae were counted per strain; note that septum formation is given relative to the values of unipolar or bipolar hyphae set to 100%). For statistical analysis, the percentage of bipolarity was investigated by using unpaired two-tailed Student′s t-test. Three independent experiments (n = 3) were conducted with at least 10 hyphae per strain. (**D**) Relative chitinase activity mainly detecting chitinase Cts1 [16] in yeast (left) or hyphal form (right, error bars, s.e.m.; n = 5 independent experiments). For statistical analysis, the relative chitinolytic activity was tested by using unpaired two-tailed Student′s t-test. Five independent experiments (n = 5) were conducted. (**E**) Western blot analysis demonstrating equal amounts of Upa2-Gfp versions using αGfp antibody (top panel). Expected molecular weight: Upa2-Gfp, 259 kDa; Pab1-Gfp, 98 kDa; Upa1-Gfp, 166 kDa. Note, that the molecular weight of Upa1-Gfp appeared larger in gel electrophoresis. Actin served as a loading control using α-actin antibody (bottom panel; expected size for actin, 42 kDa). Asterisks in αGfp panels mark putative degradation products of Gfp-tagged Upa2.

**Figure EV3.**
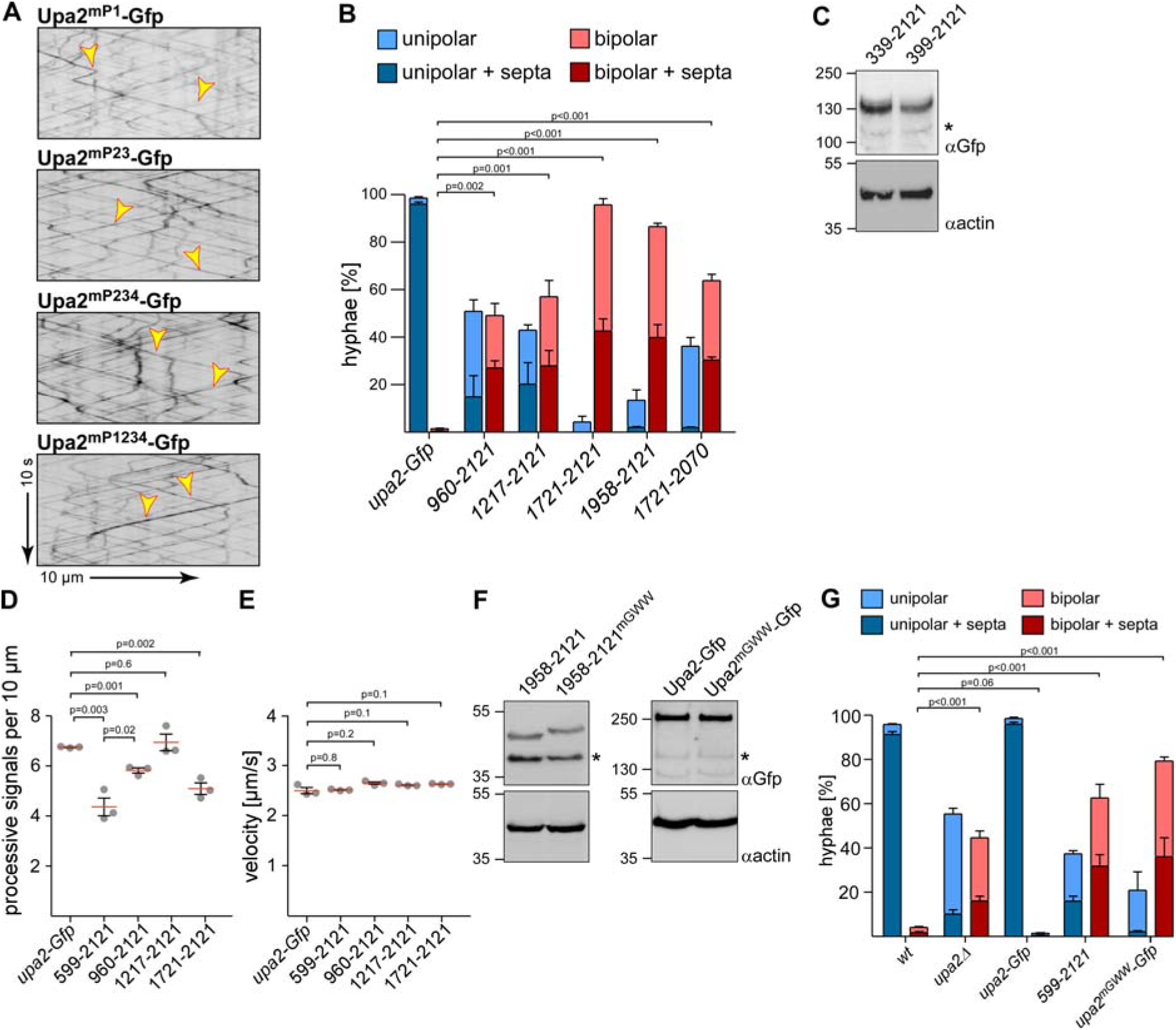
The GWW motif of Upa2 is essential for correct hyphal growth. (**A**) Kymographs of AB33 hyphae (6 h.p.i.) expressing versions of Upa2-Gfp with mutations in PAM2 motif as indicated (arrow length on the left and bottom indicate time and distance, respectively). Bidirectional movement is visible as diagonal lines (yellow arrowheads). (**B**) Quantification of hyphal growth (6 h.p.i): unipolarity, bipolarity and septum formation were quantified (error bars, s.e.m.; n = 3 independent experiments, for each experiment >100 hyphae were counted per strain; note that septum formation is given relative to the values of unipolar or bipolar hyphae set to 100%). For statistical analysis, the percentage of bipolarity was investigated by using unpaired two-tailed Student′s t-test. (**C**) Western blot analysis comparing protein amounts of functional and non-functional N-terminal truncations of Upa2-Gfp (Upa2^339-2121^-Gfp, 224 kDa; Upa2^399-2121^-Gfp, 217 kD, respectively) using αGfp antibody (top panel). Note, that the molecular weight appeared smaller in gel electrophoresis, most likely due to the use of a l-arginine buffer (see Materials and Methods). Actin served as a loading control using αactin antibody (bottom panel; expected size for actin, 42 kDa). Asterisks in αGfp panels mark putative degradation products of Gfp-tagged Upa2. (**D**) Processive signals per 10 µm of hyphal length (data points representing means from n = 3 independent experiments, with mean of means, red line, and s.e.m.; unpaired two-tailed Student′s t-test. for each experiment at least 10 hyphae were analysed per strain). (E) Velocity of fluorescent signals (velocity of tracks with > 5 µm processive movement; data points representing means from n = 3 independent experiments; with mean of means, red line, and s.e.m.; unpaired two-tailed Student′s t-test, for each experiment at least 30 signals per hypha were analysed out of 10 hyphae per strain). (**F**) Western Blot analysis comparing Upa2-Gfp versions with mutations in the C-terminal GWW motif (Upa2-Gfp, 259 kDa; Upa2^mGWW^-Gfp, 258 kDa; Upa2^1958-2121^-Gfp, 46 kDa; Upa2^1958-2121mGWW^-Gfp, 46 kDa). Note that due to the change of two tryptophans to alanine, the running behaviour of Upa2^1958-2121mGWW^-Gfp was different from Upa2^1958-2121^-Gfp. Actin served as a loading control using αactin antibody (bottom panel; expected size for actin, 42 kDa). Asterisks in αGfp panels mark putative degradation products of Gfp-tagged Upa2. (**G**) Quantification of hyphal growth (6 h.p.i): unipolarity, bipolarity and septum formation were quantified (error bars, s.e.m.; n = 3 independent experiments, for each experiment >100 hyphae were counted per strain; note that septum formation is given relative to the values of unipolar or bipolar hyphae set to 100%). For statistical analysis, the percentage of bipolarity was investigated by using unpaired two-tailed Student′s t-test.

**Figure EV4.**
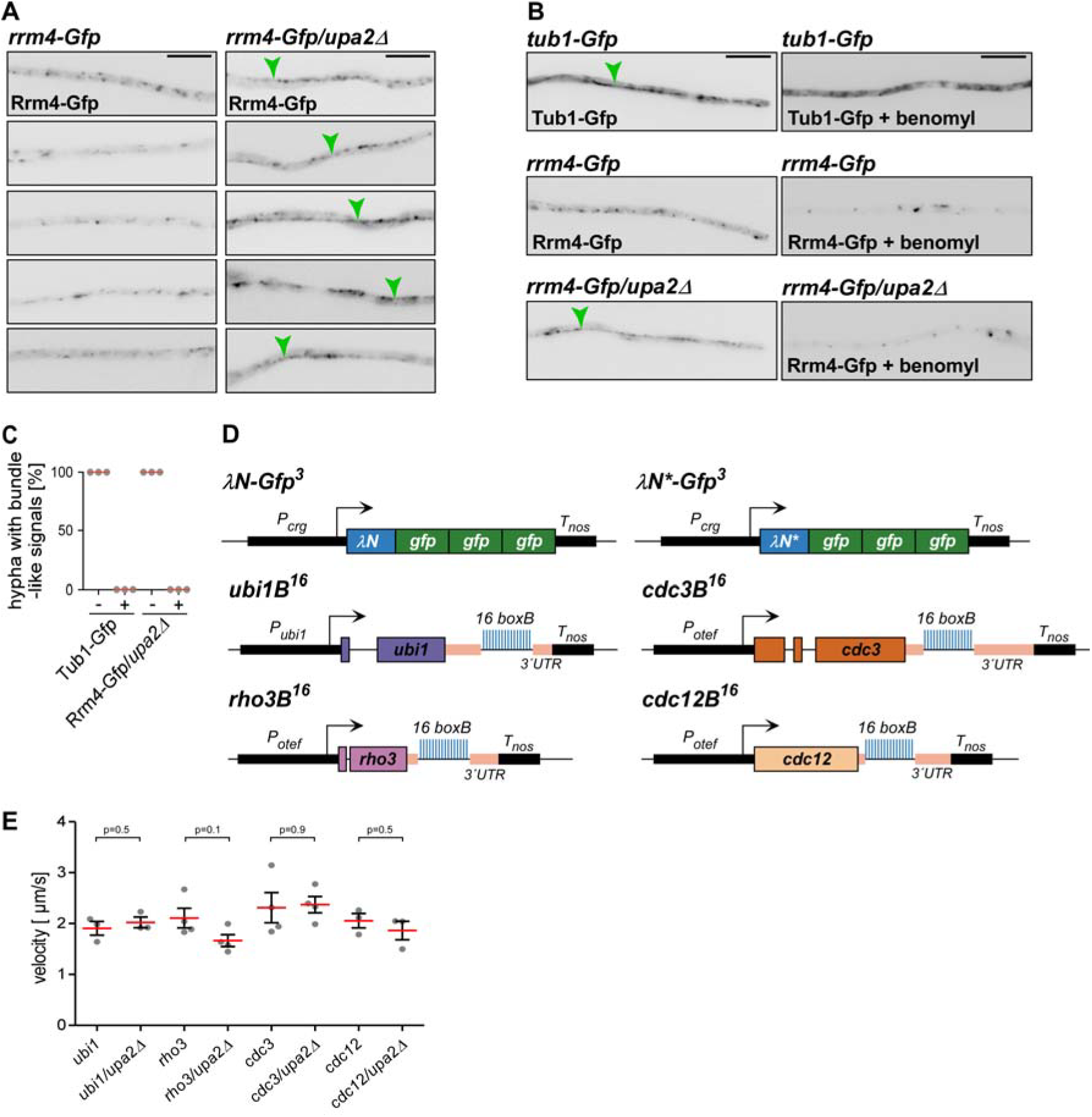
Loss of Upa2 causes aberrant bundle-like accumulation of Rrm4-Gfp. **(A)** Micrographs (inverted fluorescence image; size bar, 10 µm of AB33 hyphae (6 h.p.i.) expressing Rrm4-Gfp (green arrowheads indicate bundle-like accumulation of Rrm4-Gfp). **(B)** Micrographs (inverted fluorescence image; size bar, 10 µm) of AB33 hyphae (6 h.p.i.) expressing Tub1-Gfp (ectopic expression of Tub1 fused N-terminally with Gfp) [52] or Rrm4-Gfp (green arrowheads indicate bundle-like accumulation of Rrm4-Gfp). Hyphae on the right were treated with the microtubule inhibitor benomyl. (**C**) Quantification of bundle structures in strains expressing Gfp-Tub1 or Rrm4-Gfp with and without treatment of microtubule inhibitor benomyl (data points representing percentages from n = 3 independent experiments, with mean of means, red line; for each experiment at least 10 hyphae were analysed per strain). (**D**) (**A**) Schematic representation of the λN / λN* RNA live imaging system (P_otef_, constitutively active promoter; P_crg_ arabinose-induced promoter) T_nos_, heterologous transcriptional terminator; ubi1, ubiquitin fusion protein; *rho3*, small GTPase; cdc3 and cdc12, septins). All tested target genes carried 16 copies of the boxB hairpin in their 3′ UTR. λN*Gfp^3^ is recruited to mRNAs containing the λN-binding sites designated boxB [14]. (**E**) Velocity of fluorescent signals of analysed mRNAs (velocity of tracks with > 3 µm processive movement; data points representing means from n = 3 independent experiments, with mean of means, red line, and s.e.m.; unpaired two-tailed Student′s t-test; for each experiment more than seven hyphae were analysed per strain). Merged datasets of all values are shown.

**Figure EV5.**
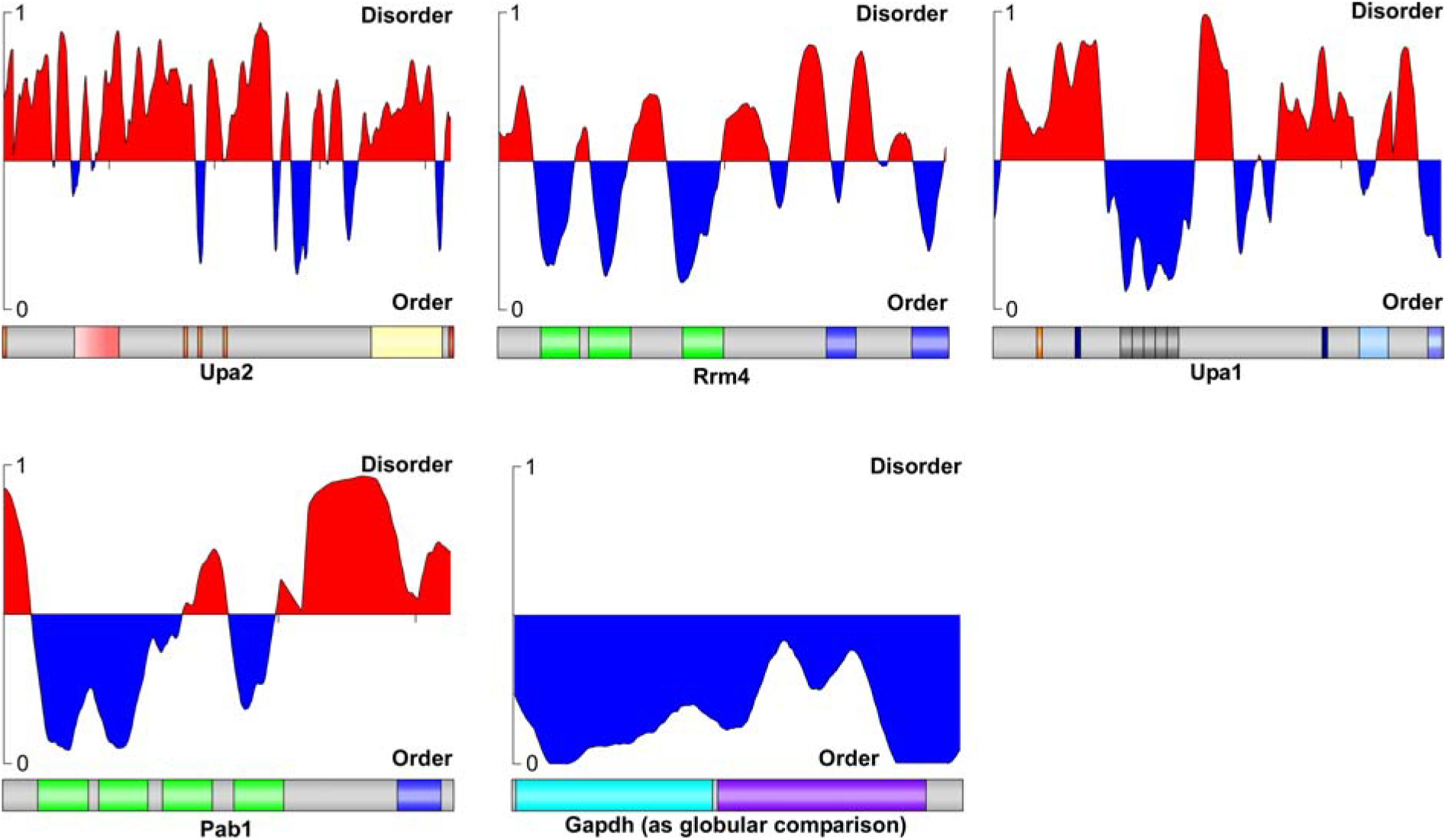
Upa2 has extensive intrinsically disordered regions. Analysis of intrinsically disordered regions using the PONDR algorithm VL3-BA. Amino acids are given prediction values between 0 and 1. Regions with values of 0.5 or above are considered disordered and are marked red, while peptide regions with values lower than 0.5 are considered ordered and are marked blue. The graphical output for the prediction is depicted in relation to the protein, shown below the graph. The size of the protein models is adjusted to the graph size. Extensive regions of Upa2 (>80%) are predicted to be disordered. Additionally, further known components of endosomal mRNA-transport, Rrm4, Upa1 and Pab1, show significant disordered stretches. Gapdh of *U. maydis* is shown as a structured protein in comparison.

