## Supplementary Information for "The multi PAM2 protein Upa2 functions as novel core component of endosomal mRNA transport"

[†] equal contribution

#### Appendix Table S1: Description of *U. maydis* strains used in this study

| | Locus | Progenitor strain | Short description |
|---|---|---|---|
| AB33 | *b* | FB2 | *Pnar:bW2bE1*, expression of active b heterodimer under control of the *nar1* promoter, strain grows filamentously upon changing the nitrogen source. |
| AB33rrm4Δ | *rrm4* | AB33 | carrying a deletion of *rrm4*. |
| AB33upa1Δ | *upa1* | AB33 | carrying a deletion of *upa1*. |
| AB33upa2Δ | *upa2* | AB33 | carrying a deletion of *upa2*. |
| AB33upa2-Gfp | *upa2* | AB33 | expressing Upa2 C-terminally fused to eGfp. |
| AB33upa2$^{mP1}$-Gfp | *upa2* | AB33upa2Δ | expressing Upa2$^{mP1}$ C-terminally fused to eGfp. Upa2$^{mP1}$ carries the amino acid substitutions L6A, V8A and F13A in the first PAM2-motif. |
| AB33upa2$^{mP234}$-Gfp | *upa2* | AB33upa2Δ | expressing Upa2$^{mP234}$ C-terminally fused to eGfp. Upa2$^{mP234}$ carries the amino acid substitutions L863A and F870A in the second PAM2-motif, the amino acid substitutions L925A und F932A in the third PAM2-motif and L1051A und F1058A in the fourth PAM2-motif. |
| AB33upa2$^{mP1234}$-Gfp | *upa2* | AB33upa2Δ | expressing Upa2$^{mP234}$ C-terminally fused to eGfp. Upa2$^{mP1234}$ carries the amino acid substitutions L6A, V8A and F13A in the first PAM2-motif, L863A and F870A in the second PAM2-motif, L925A und F932A in the third PAM2-motif and L1051A und F1058A in the fourth PAM2-motif. |
| AB33rrm4-Gfp | *rrm4* | AB33rrm4Δ | expressing Rrm4 C-terminally fused to eGfp. |
| AB33pab1-Gfp | *pab1* | AB33 | expressing Pab1 C-terminally fused to eGfp. |
| AB33upa2-Gfp/rrm4-mCherry | *upa2* *rrm4* | AB33upa2-Gfp | expressing Upa2 C-terminally fused to eGfp and Rrm4 C-terminally fused to mCherry, |
| AB33upa2-Gfp/pab1-mCherry | *upa2* *pab1* | AB33upa2-Gfp | expressing Upa2 C-terminally fused to eGfp and Pab1 C-terminally fused to mCherry, |
| AB33pab1-Gfp/rrm4Δ | *pab1* *rrm4* | AB33pab1-Gfp | expressing Pab1 C-terminally fused to eGfp and carrying a deletion of *rrm4* |
| AB33upa1-Gfp/rrm4Δ | *upa1* *rrm4* | AB33upa1-Gfp | expressing Upa1 C-terminally fused to eGfp and carrying a deletion of *rrm4* |
| AB33upa2-Gfp/rrm4Δ | *upa2* *rrm4* | AB33upa2-Gfp | expressing Upa2 C-terminally fused to eGfp and carrying a deletion of *rrm4* |
| AB33upa2-Gfp/rrm4$^{mR1}$-Rfp | *upa2* *rrm4* | AB33upa2-Gfp | expressing Upa2 C-terminally fused to eGfp and Rrm4 containing mutations in the first RRM C-terminally fused to mRFP. The conserved amino acids TVEF (aa 116-119) in the RNP2-motif of RRM1 were replaced by AAAA . |
| AB33pab1-Gfp/rrm4$^{mR1}$-Rfp | *pab1* *rrm4* | AB33pab1-Gfp | expressing Pab1 C-terminally fused to eGfp and Rrm4 containing mutations in the first RRM C-terminally fused to mRFP. The conserved amino acids TVEF (aa 116-119) in the RNP2-motif of RRM1 were replaced by AAAA . |
| AB33upa2$^{339\text{-}2121}$-Gfp | *upa2* | AB33upa2Δ | expressing Upa2$^{339–2121}$ C-terminally fused to eGfp. Like Upa2-Gfp, but carrying a N-terminal truncation from aa 1-338. |
| AB33upa2$^{399\text{-}2121}$-Gfp | *upa2* | AB33upa2Δ | expressing Upa2$^{399–2121}$ C-terminally fused to eGfp. Like Upa2-Gfp, but carrying a N-terminal truncation from aa 1-398. |
| AB33upa2$^{496\text{-}2121}$-Gfp | *upa2* | AB33upa2Δ | expressing Upa2$^{496–2121}$ C-terminally fused to eGfp. Like Upa2-Gfp, but carrying a N-terminal truncation from aa 1-495. |
| AB33upa2$^{599\text{-}2121}$-Gfp | *upa2* | AB33upa2Δ | expressing Upa2$^{599–2121}$ C-terminally fused to eGfp. Like Upa2-Gfp, but carrying a N-terminal truncation from aa 1-598. |
| AB33upa2$^{960\text{-}2121}$-Gfp | *upa2* | AB33upa2Δ | expressing Upa2$^{960–2121}$ C-terminally fused to eGfp. Like Upa2-Gfp, but carrying a N-terminal truncation from aa 1-959. |
| AB33upa2$^{1217\text{-}2121}$-Gfp | *upa2* | AB33upa2Δ | expressing Upa2$^{1217–2121}$ C-terminally fused to eGfp. Like Upa2-Gfp, but carrying a N-terminal truncation from aa 1-1216. |
| AB33upa2$^{1721\text{-}2121}$-Gfp | *upa2* | AB33upa2Δ | expressing Upa2$^{1712–2121}$ C-terminally fused to eGfp. Like Upa2-Gfp, but carrying a N-terminal truncation from aa 1-1720. |

| | | | |
|---|---|---|---|
| AB33upa2$^{1958\text{-}2121}$-Gfp | *upa2* | AB33upa2Δ | expressing Upa2$^{1958\text{–}2121}$ C-terminally fused to eGfp. Like Upa2-Gfp, but carrying a N-terminal truncation from aa 1-1957. |
| AB33upa2$^{1721\text{-}2070}$-Gfp | *upa2* | AB33upa2Δ | expressing Upa2$^{1721\text{–}2070}$ C-terminally fused to eGfp. Like Upa2-Gfp, but carrying a N-terminal truncation from aa 1-1720 and a C-terminal truncation from 2071-2121. |
| AB33upa2$^{1958\text{-}2121_mGWW}$-Gfp | *upa2* | AB33upa2Δ | expressing Upa2$^{1958\text{-}2121_mGWW}$ C-terminally fused to eGfp. Like Upa2$^{1958\text{-}2121}$-Gfp, but carrying the amino acid substitutions G2118A, W2019A and W2120A in the GWW-motif. |
| AB33upa2$^{mGWW}$-Gfp | *upa2* | AB33upa2Δ | expressing Upa2$^{mGWW}$ C-terminally fused to eGfp. Like Upa2-Gfp, but carrying the amino acid substitutions G2118A, W2019A and W2120A in the GWW-motif. |
| AB33rrm4-Gfp/upa2Δ | *rrm4*<br>*upa2* | AB33rrm4-Gfp | expressing Rrm4 C-terminally fused to eGfp and carrying a deletion of *upa2*. |
| AB33pab1-Gfp/upa2Δ | *pab1*<br>*upa2* | AB33pab1-Gfp | expressing Pab1 C-terminally fused to eGfp and carrying a deletion of *upa2*. |
| AB33cdc3B$^{16}$/λN*-Gfp$^{3}$ | *cdc3*<br>*ip$^S$* | AB33cdc3B$^{16}$ | expressing λN*-peptide fused to triple eGfp and *cdc3* mRNA tagged with 16 copies of boxB binding site. *cdc3* mRNA is under the control of the constitutive *otef* promoter. |
| AB33cdc3B$^{16}$/λN*-Gfp$^{3}$/upa2Δ | *cdc3*<br>*ip$^S$*<br>*upa2* | AB33cdc3B$^{16}$/λN*-Gfp$^{3}$ | expressing and λN*-peptide fused to triple eGfp and *cdc3* mRNA tagged with 16 copies of boxB binding site. *cdc3* mRNA is under the control of the constitutive *otef* promoter. Additionally carrying a deletion of *upa2*. |
| AB33ubi1B$^{16}$/λN-Gfp$^{3}$ | *cdc3*<br>*ip$^S$* | AB33ubi1B$^{16}$ | expressing λN-peptide fused to triple eGfp and *ubi1* mRNA tagged with 16 copies of boxB binding site. *ubi1* mRNA is under control by native promoter. |
| AB33ubi1B$^{16}$/λN-Gfp$^{3}$/upa2Δ | *cdc3*<br>*ip$^S$*<br>*upa2* | AB33ubi1B$^{16}$/λN*-Gfp$^{3}$ | expressing and λN-peptide fused to triple eGfp and *ubi1* mRNA tagged with 16 copies of boxB binding site. *ubi1* mRNA is under control by native promoter. Additionally carrying a deletion of *upa2*. |
| AB33rho3B$^{16}$/λN-Gfp$^{3}$ | *cdc3*<br>*ip$^S$* | AB33rho3B$^{16}$ | expressing λN-peptide fused to triple eGfp and *rho3* mRNA tagged with 16 copies of boxB binding site. *rho3* mRNA is under control by by native promoter. |
| AB33rhoB$^{16}$/λN-Gfp$^{3}$/upa2Δ | *cdc3*<br>*ip$^S$*<br>*upa2* | AB33rho3B$^{16}$/λN*-Gfp$^{3}$ | expressing and λN-peptide fused to triple eGfp and *rho3* mRNA tagged with 16 copies of boxB binding site. *rho3* mRNA is under control by native promoter. Additionally carrying a deletion of *upa2*. |
| AB33cdc12B$^{16}$/λN*-Gfp$^{3}$ | *cdc3*<br>*ip$^S$* | AB33cdc12B$^{16}$ | expressing λN*-peptide fused to triple eGfp and *cdc12* mRNA tagged with 16 copies of boxB binding site. *cdc12* mRNA is under the control of the constitutive *otef* promoter |
| AB33cdc12B$^{16}$/λN*-Gfp$^{3}$/upa2Δ | *cdc3*<br>*ip$^S$*<br>*upa2* | AB33cdc12B$^{16}$/λN*-Gfp$^{3}$ | expressing and λN*-peptide fused to triple eGfp and *cdc12* mRNA tagged with 16 copies of boxB binding site. *cdc12* mRNA is under the control of the constitutive *otef* promoter. Additionally carrying a deletion of *upa2*. |
| AB33cdc3-Gfp | *cdc3* | AB33 | expressing Cdc3 N-terminally fused to eGfp (see Supplementary Figure 1 in Zander, 2016). |
| AB33cdc3-Gfp/upa2Δ | *cdc3*<br>*upa2* | AB33cdc3-Gfp | expressing Cdc3 N-terminally fused to eGfp and carrying a deletion of *upa2*. |
| AB33tub1-Gfp | *tub1* | AB33 | expressing Tub1 (alpha tubulin) N-terminally fused to eGfp. *Tub1-Gfp* is inserted ectopically in the *ip$^S$* locus and under the control of the constitutive *otef* promoter |

**Appendix Table S2: Generation of *U. maydis* strains used in this study**

Uma and pUMa, internal reference numbers for strains and plasmids, respectively.

| Strains | Relevant genotype | Uma | Reference | Transformed plasmid | Locus | Progenitor |
|---|---|---|---|---|---|---|
| AB33 | *a2 $P_{nar}$:bW2 bE1* | 133 | Brachmann, 2001 | pAB33 | *b* | FB2 |
| AB33rrm4Δ | *rrm4Δ* | 273 | Becht, 2006 | pRrm4Δ_HygR (pUMa495) | *rrm4* | AB33 |
| AB33upa1Δ | *upa1Δ* | 859 | Pohlmann, 2015 | pUpa1Δ_HygR (pUMa1574) | *upa1* | AB33 |
| AB33upa2Δ | *upa2Δ* | 1505 | this study | pUpa2Δ_NatR (pUMa2408) | *upa2* | AB33upa2Δ_HygR |
| AB33upa2-Gfp | *upa2-Gfp* | 1791 | this study | pUpa2-Gfp _HygR (pUMa2654) | *upa2* | AB33upa2Δ |
| AB33upa2$^{mP1}$-Gfp | *upa2$^{mP1}$-Gfp* | 2373 | this study | pUpa1$^{mP1}$-Gfp _NatR (pUMa3422) | *upa2* | AB33upa2Δ |
| AB33upa2$^{mP234}$-Gfp | *upa2$^{mP234}$-Gfp* | 2375 | this study | pUpa1$^{mP234}$-Gfp _NatR (pUMa3424) | *upa2* | AB33upa2Δ |
| AB33upa2$^{mP1234}$-Gfp | *upa2$^{mP1234}$-Gfp* | 2376 | this study | pUpa1$^{mP1234}$-Gfp _NatR (pUMa3426) | *upa2* | AB33upa2Δ |
| AB33rrm4-Gfp | *rrm4-Gfp* | 274 | Becht, 2006 | pRrm4-Gfp_NatR (pUMa496) | *rrm4* | AB33rrm4Δ |
| AB33pab1-Gfp | *pab1-Gfp* | 389 | König, 2009 | pPab1-Gfp_NatR (pUMa805) | *pab1* | AB33 |
| AB33upa2-Gfp/rrm4-mCherry | *rrm4-mCherry* | 2132 | this study | pRrm4-mCherry _NatR (pUMa2964) | *rrm4* | AB33upa2-Gfp |
| AB33upa2-Gfp/pab1-mCherry | *rab1-mCherry* | 2131 | this study | pPab1-mCherry _NatR (pUMa2963) | *pab1* | AB33upa2-Gfp |
| AB33pab1-Gfp/rrm4Δ | *rrm4Δ* | 472 | König, 2009 | pRrm4Δ_HygR (pUMa495) | *rrm4* | AB33pab1-Gfp |
| AB33upa1-Gfp/rrm4Δ | *rrm4Δ* | 1048 | Pohlmann, 2015 | pRrm4Δ2_HygR (pUMa1391) | *rrm4* | AB33upa1-Gfp |
| AB33upa2-Gfp/rrm4Δ | *rrm4Δ* | 515 | this study | pRrm4Δ_HygR (pUMa495) | *rrm4* | AB33upa2-Gfp |
| AB33upa2-Gfp/rrm4$^{mR1}$-Rfp | *upa2-Gfp rrm4$^{mR1}$-Rfp* | 896 | this study | pRrm4$^{mR1}$-Rfp_HygR (pUMa1006) | *rrm4* | AB33upa2-Gfp |
| AB33pab1-Gfp/rrm4$^{mR1}$-Rfp | *pab1-Gfp rrm4$^{mR1}$-Rfp* | 758 | Baumann, 2014 | pRrm4$^{mR1}$-Rfp_HygR (pUMa1006) | *rrm4* | AB33pab1-Gfp |
| AB33upa2$^{339\text{-}2121}$-Gfp | *upa2$^{339\text{-}2121}$-Gfp* | 2715 | this study | pUpa2$^{339\text{-}2121}$-Gfp _HygR (pUMa3897) | *upa2* | AB33upa2Δ |

| | | | | | | |
|---|---|---|---|---|---|---|
| AB33upa2$^{399-2121}$-Gfp | *upa2$^{399-2121}$-Gfp* | 2613 | this study | pUpa2$^{399-2121}$-Gfp _HygR (pUMa3719) | *upa2* | AB33upa2Δ |
| AB33upa2$^{496-2121}$-Gfp | *upa2$^{496-2121}$-Gfp* | 2615 | this study | pUpa2$^{496-2121}$-Gfp _HygR (pUMa3720) | *upa2* | AB33upa2Δ |
| AB33upa2$^{599-2121}$-Gfp | *upa2$^{599-2121}$-Gfp* | 1789 | this study | pUpa2$^{599-2121}$-Gfp _HygR (pUMa2652) | *upa2* | AB33upa2Δ |
| AB33upa2$^{960-2121}$-Gfp | *upa2$^{960-2121}$-Gfp* | 1788 | this study | pUpa2$^{960-2121}$-Gfp _HygR (pUMa2651) | *upa2* | AB33upa2Δ |
| AB33upa2$^{1217-2121}$-Gfp | *upa2$^{1217-2121}$-Gfp* | 1787 | this study | pUpa2$^{1217-2121}$-Gfp _HygR (pUMa2650) | *upa2* | AB33upa2Δ |
| AB33upa2$^{1721-2121}$-Gfp | *upa2$^{1721-2121}$-Gfp* | 1493 | this study | pUpa2$^{1721-2121}$-Gfp _HygR (pUMa2394) | *upa2* | AB33upa2Δ |
| AB33upa2$^{1958-2121}$-Gfp | *upa2$^{1958-2121}$-Gfp* | 1763 | this study | pUpa2$^{1958-2121}$-Gfp _HygR (pUMa2721) | *upa2* | AB33upa2Δ |
| AB33upa2$^{1721-2070}$-Gfp | *upa2$^{1721-2070}$-Gfp* | 1476 | this study | pUpa2$^{1721-2070}$-Gfp _HygR (pUMa2432) | *upa2* | AB33upa2Δ |
| AB33upa2$^{1958-2121_mGWW}$-Gfp | *upa2$^{1958-2121_mGWW1}$-Gfp* | 2176 | this study | pUpa2$^{1958-2121_mGWW}$-Gfp _HygR (pUMa3180) | *upa2* | AB33upa2Δ |
| AB33upa2$^{mGWW}$-Gfp | *upa2$^{mGWW}$-Gfp* | 1893 | this study | pUpa2$^{mGWW}$-Gfp _HygR (pUMa2862) | *upa2* | AB33upa2Δ |
| AB33rrm4-Gfp/upa2Δ | *rrm4-Gfp upa2Δ* | 1164 | this study | pUpa2Δ_HygR (pUMa691) | *upa2* | AB33rrm4-Gfp |
| AB33pab1-Gfp/upa2Δ | *pab1-Gfp upa2Δ* | 1165 | this study | pUpa2Δ_HygR (pUMa691) | *upa2* | AB33pab1-Gfp |
| AB33cdc3B$^{16}$/λN*-Gfp$^{3}$ | *cdc3B$^{16}$ λN*-Gfp$^{3}$* | 848 | Baumann, 2014 | pP$_{otef}$cdc3B$^{16}$-3‘UTR-NatR (pUMa1448) | *ip$^{S}$* | AB33*λN*-Gfp$^{36}$* |
| AB33cdc3B$^{16}$/λN*-Gfp$^{3}$ /upa2Δ | *cdc3B$^{16}$ λN*-Gfp$^{3}$ upa2Δ* | 2887 | this study | pUpa2Δ_HygR (pUMa691) | *upa1* | AB33cdc3B$^{16}$ /λN*-Gfp$^{3}$ |
| AB33ubi1B$^{16}$/λN-Gfp$^{3}$ | *ubi1B$^{16}$ λN-Gfp$^{3}$* | 415 | König, 2009 | pP$_{crg1}$λN-Gfp _CbxR (pUMa975) | *ip$^{S}$* | AB33ubi1B$^{16}$ |
| AB33ubi1B$^{16}$/λN-Gfp$^{3}$/upa2Δ | *ubi1B$^{16}$ λN-Gfp$^{3}$ upa2Δ* | 2885 | this study | pUpa2Δ_HygR (pUMa691) | *upa1* | AB33ubi1B$^{16}$ /λN-Gfp$^{3}$ |
| AB33rho3B$^{16}$/λN-Gfp$^{3}$ | *rho3B$^{16}$ λN-Gfp$^{3}$* | 430 | König, 2009 | pP$_{otef}$rho3B$^{16}$-3‘UTR-NatR (pUMa989) | *ip$^{S}$* | AB33*λN-Gfp$^{3}$* |
| AB33rho3B$^{16}$/λN-Gfp$^{3}$/upa2Δ | *rho3B$^{16}$ λN-Gfp$^{3}$ upa2Δ* | 2886 | this study | pUpa2Δ_HygR (pUMa691) | *upa1* | AB33rho3B$^{16}$ /λN-Gfp$^{3}$ |

| | | | | | | |
|---|---|---|---|---|---|---|
| AB33cdc12B$^{16}$/λN*-Gfp$^{3}$ | *cdc12B$^{16}$ λN*-Gfp$^{3}$* | 849 | Zander, 2016 | pP$_{otef}$cdc12B$^{16}$-3'UTR-NatR (pUMa1449) | *ip$^{S}$* | AB33*λN*-Gfp$^{3}$* |
| AB33cdc12B$^{16}$/λN*-Gfp$^{3}$/upa2Δ | *cdc12B$^{16}$ λN*-Gfp$^{3}$ upa2Δ* | 2888 | this study | pUpa2Δ_HygR (pUMa691) | *upa1* | AB33cdc12B$^{16}$ /λN*-Gfp$^{3}$ |
| AB33cdc3-Gfp | *cdc3-Gfp* | 449 | Baumann, 2014 | pCdc3-Gfp$^{N}$_NatR (pUMa1028) | *cdc3* | AB33 |
| AB33cdc3-Gfp/upa2Δ | *cdc3-Gfp upa2Δ* | 941 | this study | pUpa2Δ_HygR (pUMa691) | *upa1* | AB33cdc3-Gfp |
| AB33tub1-Gfp | *tub1-Gfp* | 498 | This study | pTub1-Gfp_CbxR (pUMa508) | *tub1* | AB33 |

**Appendix Table S3: Description of plasmids used for *U. maydis* strain generation**
Uma and pUMa, internal reference numbers for strains and plasmids, respectively.

| Plasmid | pUMa | Resistance cassette | Short description |
|---|---|---|---|
| pAB33 | 705 | SfiI-insert of pMF2-2p | Published (Brachmann, 2001) |
| pRrm4Δ_HygR | 495 | SfiI-insert of pMF1hs | Published (Becht, 2006) |
| pUpa1Δ_HygR | 1574 | SfiI-insert of pMF1hs | Published (Pohlmann, 2015) |
| pUpa2Δ_NatR | 2408 | SfiI-insert of pMF1n | Plasmid for generating deletion mutants of *upa2*. Resistance cassette is flanked by 2 kb upstream and 1.4 kb downstream region of *upa2*. Flanking regions were amplified by PCR using oMF939/oMF940 and oMF941/oMF942 and UM521 wild-type DNA as template. |
| pUpa2-Gfp _HygR | 2654 | SfiI-insert of pMF5-1h (with removed NdeI) | Plasmid for generating eGfp fusions of *upa2*. A cassette containing Gfp, the $T_{nos}$ terminator and a hygromycin resistance cassette is flanked by 7.5 kb upstream region including 6.5 kb of the *upa1* ORF and 1.2 kb downstream region of *upa1*. Flanking regions were amplified by PCR using oDD160/oDD161 and oDD162/oDD163 and UM521 wild-type DNA as template. |
| pUpa2$^{mP1}$-Gfp _NatR | 3422 | SfiI-insert of pMF5-1n (with removed NdeI) | Plasmid for generating eGfp fusions of *upa2*$^{mP1}$. Like pUpa2-Gfp_NatR, but carrying the amino acid substitutions L6A, V8A and F13A in the first PAM2-motif. |
| pUpa2$^{mP234}$-Gfp _NatR | 3424 | SfiI-insert of pMF5-1n (with removed NdeI) | Plasmid for generating eGfp fusions of *upa2*$^{mP234}$. Like pUpa2-Gfp_NatR, but carrying the amino acid substitutions L863A and F870A in the second PAM2-motif, the amino acid substitutions L925A und F932A in the third PAM2-motif and L1051A und F1058A in the fourth PAM2-motif. |
| pUpa2$^{mP1234}$-Gfp _NatR | 3426 | SfiI-insert of pMF5-1n (with removed NdeI) | Plasmid for generating eGfp fusions of *upa2*$^{mP1234}$. Like pUpa2-Gfp_NatR, but carrying the amino acid substitutions L6A, V8A and F13A in the first PAM2-motif, the amino acid substitutions L863A and F870A in the second PAM2-motif, the amino acid substitutions L925A und F932A in the third PAM2-motif and L1051A und F1058A in the fourth PAM2-motif. |
| pRrm4-Gfp_NatR | 496 | SfiI-insert of pMF5-1n | Published (Becht, 2006). |
| pPab1-Gfp_NatR | 805 | SfiI-insert of pMF5-1n | Published (König, 2009). |
| pRrm4-mCherry _NatR | 2964 | SfiI-insert of pMF5-5n | Plasmid for generating mCherry fusions of *rrm4*. Like pRrm4-Gfp_NatR, but the C-terminal Gfp-tag was exchanged with an mCherry-tag. |
| pPab1-mCherry _NatR | 2963 | SfiI-insert of pMF5-5n | Plasmid for generating mCherry fusions of *pab1*. Like pPab1-Gfp_NatR, but the C-terminal Gfp-tag was exchanged with an mCherry-tag. |
| pRrm4Δ2_HygR | 1391 | SfiI-insert of pMF1hl | Published (Pohlmann, 2015) |
| pRrm4$^{mR1}$-Rfp_HygR | 1006 | SfiI-insert of pMF5-2h | Published (Baumann, 2014) |
| pUpa2$^{284\text{-}2121}$-Gfp_HygR | 2653 | SfiI-insert of pMF5-1h (with removed NdeI) | Plasmid for generating eGfp fusions of *upa2*$^{284\text{-}2121}$. Like pUpa2-Gfp_NatR, but carrying a N-terminal truncation from aa 1-283. |
| pUpa2$^{339\text{-}2121}$-Gfp_HygR | 3897 | SfiI-insert of pMF5-1h (with removed NdeI) | Plasmid for generating eGfp fusions of *upa2*$^{339\text{-}2121}$. Like pUpa2-Gfp_NatR, but carrying a N-terminal truncation from aa 1-338. |
| pUpa2$^{399\text{-}2121}$-Gfp_HygR | 3719 | SfiI-insert of pMF5-1h (with removed NdeI) | Plasmid for generating eGfp fusions of *upa2*$^{399\text{-}2121}$. Like pUpa2-Gfp_NatR, but carrying a N-terminal truncation from aa 1-398. |
| pUpa2$^{496\text{-}2121}$-Gfp_HygR | 3720 | SfiI-insert of pMF5-1h (with removed NdeI) | Plasmid for generating eGfp fusions of *upa2*$^{496\text{-}2121}$. Like pUpa2-Gfp_NatR, but carrying a N-terminal truncation from aa 1-495. |
| pUpa2$^{599\text{-}2121}$-Gfp_HygR | 2652 | SfiI-insert of pMF5-1h (with removed NdeI) | Plasmid for generating eGfp fusions of *upa2*$^{599\text{-}2121}$. Like pUpa2-Gfp_NatR, but carrying a N-terminal truncation from aa 1-598. |
| pUpa2$^{960\text{-}2121}$-Gfp_HygR | 2651 | SfiI-insert of pMF5-1h (with removed NdeI) | Plasmid for generating eGfp fusions of *upa2*$^{960\text{-}2121}$. Like pUpa2-Gfp_NatR, but carrying a N-terminal truncation from aa 1-959. |
| pUpa2$^{1217\text{-}2121}$-Gfp_HygR | 2650 | SfiI-insert of pMF5-1h (with removed NdeI) | Plasmid for generating eGfp fusions of *upa2*$^{1217\text{-}2121}$. Like pUpa2-Gfp_NatR, but carrying a N-terminal truncation from aa 1-1216. |

| | | | |
|---|---|---|---|
| pUpa2$^{1721-2121}$-Gfp_HygR | 2394 | SfiI-insert of pMF5-1h (with removed NdeI) | Plasmid for generating eGfp fusions of *upa2$^{1721-2121}$*. Like pUpa2-Gfp_NatR, but carrying a N-terminal truncation from aa 1-1720. |
| pUpa2$^{1958-2121}$-Gfp_HygR | 2721 | SfiI-insert of pMF5-1h (with removed NdeI) | Plasmid for generating eGfp fusions of *upa2$^{1958-2121}$*. Like pUpa2-Gfp_NatR, but carrying a N-terminal truncation from aa 1-1957. |
| pUpa2$^{1721-2070}$-Gfp_HygR | 2432 | SfiI-insert of pMF5-1h (with removed NdeI) | Plasmid for generating eGfp fusions of *upa2$^{1721-2070}$*. Like pUpa2-Gfp_NatR, but carrying a N-terminal truncation from aa 1-1720 and a C-terminal truncation from 2071-2121. |
| pUpa2$^{1958-2121_mGWW}$-Gfp_HygR | 3180 | SfiI-insert of pMF5-1h (with removed NdeI) | Plasmid for generating eGfp fusions of *upa2$^{1958-2121_mGWW}$*. Like pUpa2$^{1958-2121}$-Gfp_NatR, but carrying the amino acid substitutions G2118A, W2019A and W2120A in the GWW-motif. |
| pUpa2$^{mGWW}$-Gfp_HygR | 2862 | SfiI-insert of pMF5-1h (with removed NdeI) | Plasmid for generating eGfp fusions of *upa2$^{mGWW1}$*. Like pUpa2-Gfp_NatR, but carrying the amino acid substitutions G2118A, W2019A and W2120A in the GWW-motif. |
| pUpa2Δ_HygR | 691 | SfiI-insert of pMF1hs | Plasmid for generating deletion mutants of *upa2*. Lie pUpa2Δ_NatR, but the Sfi-insert of pMF1n containing the nourseothricin resistance cassette has been replaced by the SfiI-insert of pMF1hs containing an hygromycin resistance cassette. |
| pP$_{otef}$cdc3B$^{16}$-3‘UTR-NatR | 1448 | SfiI-insert of pMF1n | Published (Zander, 2016) |
| pP$_{crg1}$λN-Gfp _CbxR | 975 | CbxR for integration at *ip$^{S}$* locus | Published (König, 2009) |
| pP$_{otef}$rho3B$^{16}$-3‘UTR-NatR | 989 | SfiI-insert of pMF1n | Published (König, 2009) |
| pP$_{otef}$cdc12B$^{16}$-3‘UTR-NatR | 1449 | SfiI-insert of pMF1n | Published (Zander, 2016) |
| pCdc3-Gfp$^{N}$_NatR | 1028 | SfiI-insert of pMF1n | Published (Baumann, 2014). Plasmid for generating of a N-terminal eGfp fusions of *cdc3*. 1.7 kb upstream flanking region of the *cdc3* locus, followed by the eGFP coding sequence with the stop codon removed. Downstream 1440 bps of the *cdc3* ORF containing all 3 exons and the stop codon is transcriptionally fused. This construct is flanked by 600 bps cdc3 3´ UTR. A 1.8 kb NatR cassette is placed downstream, followed by a 1.1 kb *cdc3* downstream flanking region. This construct is integrated into the native *cdc3* locus and preserves the Rrm4 binding site in the 3´UTR as well as the observed crosslinking regions in the 5´ UTR and the first exon (Olgeiser, 2019). |
| pTub1-Gfp_CbxR | 508 | CbxR for integration at *ip$^{S}$* locus | Plasmid for the extopic expression of Tub1 (alpha tubulin) N-terminally fused to eGfp. Tub1-Gfp is under control of the constitutive acitve *otef* promoter. |

### Appendix Table S4: Description of plasmids used for yeast two-hybrid analyses

| Plasmid | pUMa | Gene | Short description |
|---|---|---|---|
| pGADT7-DS | *1624* | | Plasmid for the expression of hybrid proteins, N-terminally fused to a nuclear localisation signal (NLS) of the simian virus 40 (SV40), followed by the Gal4 activation domain (aa 768-881) and an HA-epitope for Western Blot detection. Resulting hybrid proteins are termed AD-"X". For positive selection of transformants on minimal medium this plasmid carries a *LEU2* auxotrophy marker. This plasmid contains two diverse SfiI-restriction sites for cloning purposes (Dualsystems Biotech, Schlieren, Switzerland). |
| pGBKT7-SfiI MCS | *1625* | | Plasmid for the expression of hybrid proteins, N-terminally fused to the Gal4 DNA-binding domain (aa 1-147), followed by a c-Myc-epitope for Western Blot detection. Resulting hybrid proteins are termed BD-"X". For positive selection of transformants on minimal medium, this plasmid carries a *TRP1* auxotrophy marker. This plasmid contains two diverse SfiI-restriction sites for cloning purposes. (Clontech Laboratories, Inc., Mountain View, CA, USA). |
| pGADT7-T | *1636* | | Plasmid for the expression of a N-terminal AD-fusion of the large T-antigen of SV40. It interacts with BD-p53 as a positive control (Clontech). |
| pGBKT7-p53 | *1638* | | Plasmid for the expression of a N-terminal BD-fusion of the murine p53. It interacts with AD-T as a positive control (Clontech). |
| pGBKT7-Lam | *1637* | | Plasmid for the expression of a N-terminal BD-fusion with the human nuclear protein Lamin C, which shows no interaction with most proteins and serves as negative control (Clontech). |
| pGBKT7-Rrm4 | *1628* | *rrm4* | Plasmid for the expression of BD-Rrm4. |
| pGBKT7-Pab1 | *1632* | *pab1* | Plasmid for the expression of BD-Pab1. |
| pGADT7-Upa2 | *1631* | *upa2* | Plasmid for the expression of AD-Upa2. |
| pGADT7_upa2_1-1216 | *3562* | *upa2* | Plasmid for the expression of AD-Upa2_1-1216. Like AD-Upa2 but carrying a C-terminal truncation from aa 1217-2121. |
| pGADT7_upa2_1217-2121 | *3437* | *upa2* | Plasmid for the expression of AD-Upa2_1-1216. Like AD-Upa2 but carrying a N-terminal truncation from aa 1-1216. |
| pGBKT7-Pab1$^{mM}$ | *1998* | *pab1* | Plasmid for the expression of BD-Pab1$^{mM}$. Like BD-Pab1, but carrying the amino acid substitutions Y580A, V583A, K593A and I598A in the MLLE domain. |
| pGBKT7_onlyMLLEpab1 | *2000* | *pab1* | Plasmid for the expression of BD- MLLE$^{Pab1}$. For this the last 86 aa of the *pab1* ORF including the MLLE domain were fused to Gal4-BD and Myc. |
| pGADT7_upa2_1-1216$^{mP1}$ | *3563* | *upa2* | Plasmid for the expression of AD- Upa2_1-1216$^{mP1}$. Like AD-Upa2_1-1216, but carrying the amino acid substitutions L6A, V8A and F13A in the first PAM2-motif. |
| pGADT7_upa2_1-1216$^{mP234}$ | *3564* | *upa2* | Plasmid for the expression of AD- Upa2_1-1216$^{mP1}$. Like AD-Upa2_1-1216, but carrying the amino acid substitutions L863A and F870A in the second PAM2-motif, the amino acid substitutions L925A und F932A in the third PAM2-motif and L1051A und F1058A in the fourth PAM2-motif. |
| pGADT7_upa2_1-1216$^{mP1234}$ | *3565* | *upa2* | Plasmid for the expression of AD- Upa2_1-1216$^{mP1}$. Like AD-Upa2_1-1216, but carrying the amino acid substitutions L6A, V8A and F13A in the first PAM2-motif, L863A and F870A in the second PAM2-motif, L925A und F932A in the third PAM2-motif and L1051A und F1058A in the fourth PAM2-motif. |
| pGBKT7-mMLLE$^{pab1}$ | *3420* | *pab1* | Plasmid for the expression of BD- mMLLE$^{Pab1}$.Like BD-mMLLE$^{Pab}$, but carrying the amino acid substitutions Y580A, V583A, K593A and I598A in the MLLE domain. |
| pGADT7-upa1_1-194 | *2002* | *upa1* | Plasmid for the expression of AD-Upa1_1-194. Expresses Upa1 as BD-fusion, but carrying a C-terminal truncation from aa 195-1287. |
| pGADT7_upa2_1721-2121 | *3313* | *upa2* | Plasmid for the expression of AD-Upa2_1721-2121. Like AD-Upa2 but carrying a N-terminal truncation from aa 1-1720. |
| pGBKT7_upa2_1721-2121 | *3380* | *upa2* | Plasmid for the expression of BD-Upa2_1721-2121. Like AD-Upa2_1712-2121 but fused to BD. |
| pGADT7_upa2_1721-2121$^{mGWW}$ | *3392* | *upa2* | Plasmid for the expression of AD-Upa2_1721-2121$^{mGWW}$. Like AD-Upa2_1712-2121, but carrying a triple amino acid exchange of the conserved GWW to AAA (aa 2118-2120) |
| pGBKT7-Upa1 | *1730* | *upa1* | Plasmid for the expression of BD-Upa1. |
| pGADT7-Rrm4 | *1629* | *rrm4* | Plasmid for the expression of AD-Rrm4. |

#### Appendix Table S5: Description of plasmids used for pulldown experiments

| Plasmid | pUMa | Short description |
|---|---|---|
| pGEX_GST | 1881 | Plasmid for the expression of the GST (Glutathione S transferase)-tag alone. Expression is regulated by tac promoter. The plasmid also contains a *lacI$^q$* gene for use in *E. coli*. The plasmid carries an ampicillin resistance for selection. This vector is based on pGEX-2T (GE Healthcare). |
| pGEX_GST-MLLE$^{Pab1}$ | 2187 | Plasmid for the expression of the GST-MLLE$^{Pab1}$. The last 86 aa of the *pab1* ORF including the MLLE domain were N-terminally fused to a GST-tag. |
| pGEX_GST-Rrm4$^{720-792}$ | 2385 | Plasmid for the expression of the GST-Rrm4$^{720-792}$. A region of Rrm4 comprising of amino acid 720-792 was N-terminally fused to a GST-tag. |
| pET15B_Upa2_834-1216 | 3468 | Plasmid for the expression of a His$_6$_ Upa2_834-1216 in *E. coli*. Upa2 (aa 834-1216) which includes the coding regions for the PAM2 motifs 2, 3, 4 was N-terminally fused to a 6x histidine-tag. Expression is regulated by lacO. The plasmid carries an ampicillin resistance for selection. |
| pET15B_Upa2_834-1216$^{mP23}$ | 3469 | Plasmid for the expression of a His$_6$_ Upa2_834-1216$^{mP23}$ in *E. coli*. Like Upa2_834-1216, but carrying the amino acid substitutions L863A and F870A in the second PAM2 motif, the amino acid substitutions L925A und F932A in the third PAM2 motif. Expression is regulated by lacO. The plasmid carries an ampicillin resistance for selection. |
| pET15B_Upa2_834-1216$^{mP234}$ | 3470 | Plasmid for the expression of a His$_6$_ Upa2_834-1216$^{mP234}$ in *E. coli*. Like Upa2_834-1216, but carrying the amino acid substitutions L863A and F870A in the second PAM2 motif, the amino acid substitutions L925A und F932A in the third PAM2 motif and L1051A und F1058A in the fourth PAM2 motif. Expression is regulated by lacO. The plasmid carries an ampicillin resistance for selection. |

#### Appendix Table S6: DNA oligonucleotides used in this study

| Designation | Nucleotide sequence (5' --> 3') | Remarks |
|---|---|---|
| oMF938 | CTCGCGTTTCCACTTGCC | *upa2* u1 (deletion) |
| oMF939 | GCTGAGTTGACGTTGGGC | *upa2* u2 (deletion) |
| oMF940 | TTCGGCCATCTAGGCCTTGAGACTGCTGCCCTCC | *upa2* u3 (deletion) |
| oMF941 | TGAGGCCTGAGTGGCCTCGAGGTGGATGAAGGCG | *upa2* d1 (deletion) |
| oMF942 | AAGCAGCGAGTCGTACGC | *upa2* d2 (deletion) |
| oMF943 | GCCTGTTGTCGTTGAGCC | *upa2* d3 (deletion) |
| oMF944 | CACCAGCACCTAGTCAGC | *upa2* p1 (deletion) |
| oMF945 | TGAATCATACCGCCCGGC | *upa2* p2 (deletion) |
| oDD144 | GGTCTCGGATCGCATATGGATGCTCTGCACAGGAGGCTCG | *upa2* ORF 1721 fwd |
| oDD145 | GGTCTCCTGGCGCGTCTTTTGCTTTTCGACCG | *upa2* ORF 2070 rev |
| oDD160 | GGTCTCGCCTGCAATATTTGTAGTCAACTGCCTCTTTG | *upa2* u2 (fusion) |
| oDD161 | GGTCTCGGATCCTTGATGAGCTAGACAGGAAAC | *upa2* u3 (fusion) |
| oDD162 | GGTCTCCGGCCAGTGGTAATGATTCGAATGCC | *upa2* d1 (fusion) |
| oDD163 | GGTCTCCCTGCAATATTCTCGGTATGCGTTAGAATG | *upa2* d2 (fusion) |
| oDD526 | GGTCTCCTGGCAGACCACCACCCGCCTTC | *upa2* ORF 2121 rev |
| oDD578 | ATAACATATGTGGGGCGATCGCCGCACG | *upa2* 1217-1414 |
| oDD579 | CGTGCGGCGATCGCCCCATCG | *upa2* 960-1216 |
| oDD580 | CTTGACCTGCCTCAGCGCGTG | *upa2* 1415-1711 |
| oDD581 | ATAACATATGAATGCGGCGCCCAACGGG | *upa2* 599-959 |
| oDD582 | CGTTGGGCGCCGCATTCGGAC | *upa2* 284-598 |
| oDD583 | GTGCCATATGGAGGGCAGCAGTC | *upa2* 1-283 |
| oDD584 | CGCGCATATGCGTGCTAGCGAACCAGGTTC | *upa2* 284-598 |
| oDD585 | GAACCTGGTTCGCTAGCACGTG | *upa2* 1-283 |
| oDD586 | ATAACATATGGGGGCCGCGGTCAAGCCGAAT | *upa2* 1415-1711 |

| | | |
|---|---|---|
| oDD587 | CATTCGGCTTGACCGCGGCCC | *upa2* 1217-1414 |
| oDD588 | CGCGCATATGGTAACTAGTGCTGATGGTGGTG | *upa2* 960-1216 |
| oDD589 | GCACCACCATCAGCACTAGTTACTGCGATGC | *upa2* 599-959 |
| oDD604 | GGTCTCGGATCCATATGAAGGCTGAAGAGCTCGCCGAAG | *upa2* 1958-2121 |
| oDD741 | GGTCTCGGATCCATATGGCTGCCGCGGCGGCCGCCGCGGCGCGTGTTCACATCGAGGATGT | mutGWW 1958 |
| oRL422 | AGAGGTTACATGGCCAAG | Y2H of *upa2* |
| oRL591 | TGAGGCCATTACGGCCCATATGGAGGGCAGCAGTCTC | Y2H of *upa2* |
| oRL592 | CGTGGCCGAGGCGGCCGTCAAGACCACCACCCGCCTTC | Y2H of *upa2* |
| oMB43 | GGTCTCGCCTGGCTGAGGCAGGTCAAGCTGAAGCCGAG | *upa2* 1712-2121 |
| oMB44 | GGGTCTCGCGGCGCCTTCATCCACCTCGACCTCCTT | mutGWW |
| oMB45 | GGTCTCGGCCGCCGCTTCTGCCAACGCGGCCACCATGGTG | mutGWW |
| oMB46 | GGTCTCGCTGCGGCGCGCCGGCCGCTTTACTTGTACA | mutGWW |
| oMB922 | GGTCTCAGCATCTATGCTCCGGTTTCGACGCC | Y2H 283 rev |
| oMB923 | GGTCTCCGGCCAATGCGGCGCCCAACGGG | Y2H 599 fwd |
| oMB926 | GGTCTCAGCATCTAAGACCACCACCCGCCTTC | Y2H 1721-2121 |
| oMB958 | GGTCTCCGGCCATGGAGGGCAGCAGTGCCAACGCCGCGGCGCCTGTGGCTAAGCCCAGCGGCG CTGCAAAC | mutPAM2-1 |
| oMB959 | GGTCTCCGGCCGATGCTCTGCACAGGAGG | Y2H 1721-2121 |
| oMB970 | GGTCTCCAGCGTCAGGATTGGCCAGCGAGATTCCAGAAGCTC | mutPAM2-2rev |
| oMB971 | GGTCTCCCGCTAAGGAGGCCAAGTTCGGCGGGACAAGCGC | mutPAM2-2fwd |
| oMB972 | GGTCTCCCGCACCCACATTGGCGTGAGCTGCATTCGTGGG | mutPAM2-3rev |
| oMB973 | GGTCTCCTGCGGCTCCGGCCACCCCGGGCTTGTTTACCTTC | mutPAM2-3fwd |
| oMB974 | GGTCTCCGGCATCAGCTGTCGCGCGGGATTCGTGTGAGAC | mutPAM2-4rev |
| oMB975 | GGTCTCCTGCCCCATCGGCCGTGCCAACTTGGGCAAAAAG | mutPAM2-4fwd |
| oMB979 | GGTCTCAGCATCTATCGAATGGATGGGACGTGTC | Y2H 1216 rev |
| oUP276 | GGTCTCAGCATCTAAGAAGCGGCGGCGCCTTC | Y2H 1721 mutGWW |
| oUP327 | TCGCATATGGAGGGCAGCAGTGCCAAC | mutGWW |

#### Appendix Table S7: Accession numbers of phylogenetic analysis

| **Organism** | **Rrm4 ortholog** | **Upa2 ortholog** | **Upa1 ortholog** |
|---|---|---|---|
| *Ustilago maydis* | XP_011389820.1 | XP_011391917.1 | XP_011388912.1 |
| *Ustilago hordei* | CCF52027.1 | CCF50315.1 | CCF52210.1 |
| *Ustilago bromivora* | SAM82601.1 | SAM85987.1 | SAM82942.1 |
| *Sporisorium reilianum* | CBQ73718.1 | CBQ69926.1 | CBQ72642.1 |
| *Sporisorium scitamineum* | CDW98458.1 | CDU22493.1 | CDW96316.1 |
| *Moesziomyces antarcticus* | XP_014657015.1 | XP_014654742.1 | XP_014656570.1 |
| *Melanopsichium pennsylvanicum* | CDI54139.1 | CDI55987.1 | CDI53177.1 |
| *Thecaphora thlaspeos** | THTG_06061 | THTG_03355 | THTG_01672 |
| *Malassezia globosa* | n.a. | n.a. | XP_001732453.1** |
| *Cryptococcus neoformans* | XP_024512316.1*** | XP_024512946.1*** | XP_569116.1*** |
| *Armillaria ostoyae* | SJK98691.1 | SJL05310.1 | SJL02266.1 |
| *Coprinopsis cinerea* | XP_001832566.2 | XP_001828502.2 | XP_001837291.2 |
| *Punctularia strigosozonata* | XP_007384300.1 | XP_007378248.1 | XP_007382070.1 |
| *Serendipita indica* | CCA67340.1 | CCA73898.1 | CCA71703.1 / CCA71704.1 |
| *Trametes versicolor* | XP_008042363.1 | XP_008033367.1 | XP_008035292.1 |
| *Phanerochaete carnosa* | XP_007393387.1 | XP_007389854.1 | XP_007391955.1 |

* *T. thlaspeos* specific gene descriptions were used, because the genome has not been published so far.
** *M. globosa* Upa1 ortholog resembles the Pib1-like type of FYVE-proteins
*** *C. neoformans* proteins show weak or partial homology to Rrm4 and Upa1, and are considered not functional, although their functionality was so far not assessed.
